# Cell-of-origin and genetic drivers define advanced bladder cancer subtypes and potential therapeutic response in mouse models

**DOI:** 10.1101/2025.07.14.664768

**Authors:** Ester Munera-Maravilla, Mercedes Pérez-Escavy, Carolina Rubio, Cristina Segovia, Iris Lodewijk, Sandra P. Nunes, Álvaro Martín de Bernardo, Ignacio A. Reina, Esther Montesinos, Lucía Morales, Víctor G. Martínez, Mónica Martínez-Fernández, Marta Dueñas, Jesús M. Paramio, Cristian Suárez-Cabrera

## Abstract

Bladder cancer (BC) remains a major clinical challenge owing to its high recurrence, limited treatment options, and molecular heterogeneity. Despite recent therapeutic advances, prognosis remains poor, and resistance is frequent, underscoring the need for improved experimental models to study tumorigenesis and therapeutic response. A key limitation of advanced BC research is the scarcity of *in vivo* models that accurately reflect invasive disease, with even fewer capturing the complexity of metastasis. To investigate how the cell-of-origin and specific combinations of driver mutations influence in bladder tumorigenesis, we developed and characterized four genetically engineered mouse models of advanced BC by targeting two combinations of tumor suppressor genes (*Pten* and *Trp53*, or *Pten*, *Trp53*, *Rb1*, and *Rbl1*) in basal or suprabasal urothelial cells through intravesical of Cre-adenovirus delivery. Loss of the retinoblastoma family reduced cancer-specific survival and was associated with more differentiated carcinomas. In both genetic backgrounds, luminal-derived tumors developed earlier but showed fewer metastatic events. Histopathological and transcriptomic analyses revealed that these tumors resemble human basal-squamous and stroma-rich subtypes, sharing regulatory networks and activated signaling pathways with human invasive tumors. Notably, tumors lacking retinoblastoma family genes exhibited increased immune infiltration, reinforcing the value of these models for diverse preclinical applications. To overcome detection and latency limitations, we established tumor-derived cell lines and generated syngeneic graft models. These were validated as preclinical platforms, exhibiting therapeutic responses to CDK4/6 inhibition and anti-PD-L1 immunotherapy. Our findings highlight the value of these novel models for studying BC progression and evaluating emerging therapeutic strategies in immunocompetent settings.

## INTRODUCTION

Bladder cancer (BC) is the ninth most commonly diagnosed cancer worldwide, with an alarming burden on the global healthcare system. It is characterized by high recurrence rates, significant tumor progression, prolonged follow-up periods, and a notable lack of effective therapeutic options (1,2). The majority of BC cases are urothelial carcinomas, which are classified into two major categories: non-muscle invasive bladder cancer (NMIBC) and muscle-invasive bladder cancer (MIBC). NMIBC accounts for approximately 75% of cases and generally has a favourable prognosis, with a five-year survival rate of nearly 70%. In contrast, MIBC, which comprises about 25% of cases, is associated with significantly worse outcomes. Patients with regionally invasive MIBC have a five-year survival rate of approximately 40%, while those with metastatic BC (mBC) face a grim prognosis, with survival rates as low as 8% (3).

Despite decades of research, therapeutic advancements in advanced BC have been limited. Beyond immune checkpoint inhibitors (ICIs), in which only a minority of patients achieves a durable response, or antibody-drug conjugates such as enfortumab vedotin or sacituzumab govitecan, which have shown promising results in patients refractory to platinum-based chemotherapy and immunotherapy, treatment options remain scarce (4,5). These disparities underscore the urgent need to improve both our understanding and management of advanced BC, which remains a major challenge in the fight against BC.

At the molecular level, MIBC and mBC are highly complex and heterogeneous diseases characterized by significant genomic instability and a broad spectrum of genetic and epigenetic alterations. Over the years, various molecular classifications of MIBC have been proposed based on transcriptomic profiling, allowing for more precise patient stratification according to prognosis and potentially serving as biomarkers of differential therapy response, although this remains under investigation (6–8). A recent genomic and transcriptomic study of mBC has compared metastatic lesions with primary invasive tumors, assessing the impact of these molecular classifications and underscoring the clinical relevance of genomic alterations as potential therapeutic targets (9). These classifications have been instrumental in advancing our understanding of BC biology, revealing specific genomic alterations associated with distinct molecular subtypes and implicated in cancer progression and dissemination. Among these, mutations affecting genes involved in cell cycle regulation, chromatin remodeling, and receptor tyrosine kinase signaling (PI3K–mTOR and RTK–MAPK pathways), as well as variations in the tumor microenvironment (TME), have been identified as crucial players (6,7,9). Several studies have emphasized the clinical relevance of molecular stratification in advanced BC, suggesting that chemotherapy and immunotherapy efficacy may vary significantly across different subtypes (8,10).

Another major challenge in advanced BC research is the limited availability of *in vivo* models that effectively reflect invasive disease, with even fewer capable of modelling metastatic spread (11,12). This limitation hampers the ability to fully capture the complexity of human advanced BC, including its diverse molecular subtypes, and to effectively evaluate novel therapeutic strategies. Patient-derived xenograft (PDX) models or heterotopic and orthotopic xenografts— where human tumor tissue or cells are implanted into immunocompromised mice—, closely mimic the heterogeneity of human tumors, but they often fail to replicate the full complexity of the TME and metastatic progression (12,13). N-butyl-N-(4-hydroxybutyl)-nitrosamine (BBN)- induced carcinogenesis is a well-established model for BC in mice and rats, but tumor heterogeneity can limit its utility as a preclinical model and metastases are rare (12,13). Genetically modified mouse models (GEMM), on the other hand, can recapitulate key genetic alterations found in human advanced BC, allowing for the study of distinct molecular mechanisms. Strong evidence suggests that different molecular subtypes of BC may arise from distinct initiating cell populations, highlighting the importance of both genetic alterations and the cell of origin in tumor development and progression. However, the available options for urothelium-specific promoters to target defined subpopulations are limited, and most models developed to date primarily lead to hyperplasia, dysplasia, and non-invasive tumors, with only a few progressing to invasive disease and very few resulting in metastases (11,12). Therefore, there is a critical need to develop more useful advanced BC models that accurately replicate the molecular characteristics and behavior of human tumors.

We previously described an immunocompetent quadruple-knockout (QKO) GEMM, characterized by Cre recombinase-dependent inactivation of the tumor suppressor genes *Pten*, *Trp53*, and *Rb1* mice (in an *Rbl1*-null background due to functional compensation of *Rb1* loss in mouse tissues), specifically targeting basal urothelial cells through adenoviral delivery of Cre under the control of the keratin 5 promoter (*Ad5-Krt5-Cre*). We have used this model as a preclinical tool to test the combination of chromatin remodeling inhibitor or cell cycle inhibitor with cisplatin (14,15). Given that concurrent *TP53* and *RB1* alterations have been identified in human tumors associated with specific molecular subtypes and increased aggressiveness (6,10), in this study, we aimed to develop and characterize new models comparing the combined loss of *Pten*, *Trp53*, and *Rb1* in *Rbl1*-deficient mice (16,17) with the loss of only *Pten* and *Trp53* (double-knockout; DKO), where the retinoblastoma (Rb) family remained unaltered. Additionally, in both models, we targeted basal urothelial cells (using *Ad5-Krt5-Cre*) and luminal urothelial cells (employing the murine *Krt20* promoter).

## RESULTS

### Development and characterization of advanced BC mouse models: role of the retinoblastoma gene family and cell of origin in tumorigenesis

The specificity and expression pattern of the *Krt5* promoter used in this study have been extensively described and characterized in previous reports (18,19) To selectively target basal urothelial cells, we generated an adenoviral vector expressing Cre recombinase under the control of the *Krt5* promoter (*Ad5-Krt5-Cre*), which we have successfully employed in previous studies (14,15). In parallel, we designed and constructed a novel adenovirus to direct Cre expression to luminal urothelial cells by cloning a 1.5 kb region upstream of the mouse *Krt20* gene start codon, corresponding to a putative promoter element (**Supplemental Figure S1A**). The specificity of this promoter was validated both *in vitro*, using human BC cell lines with varying levels of KRT20 protein expression, and *in vivo*, using *mT/mG* reporter mice (**Figure 1, A and B, and Supplemental Figure S1, B and C**). Following intravesical administration of each adenovirus into mT/mG reporter mice, we observed Cre-mediated recombination in sporadic urothelial cells of either basal (Ad5-Krt5-Cre) or luminal (*Ad5-Krt20-Cre*) identity, depending on the vector used (**Figure 1, A and B**). These events mimicked spontaneous alterations that may lead to the development of human bladder tumors. To evaluate the efficiency and completeness of Cre-mediated recombination, we isolated tumor cells from 12 independent tumors derived from the different experimental models. Genomic PCR confirmed the deletion of all target loci in each case, and RT-qPCR analysis demonstrated a marked reduction or complete absence of transcript expression for the corresponding genes. These findings indicate that recombination occurred efficiently and uniformly across all alleles in both DKO and QKO mice, regardless of whether *Ad5-Krt5-Cre* or *Ad5-Krt20-Cre* was used for gene delivery (see below).

**Figure 1.**
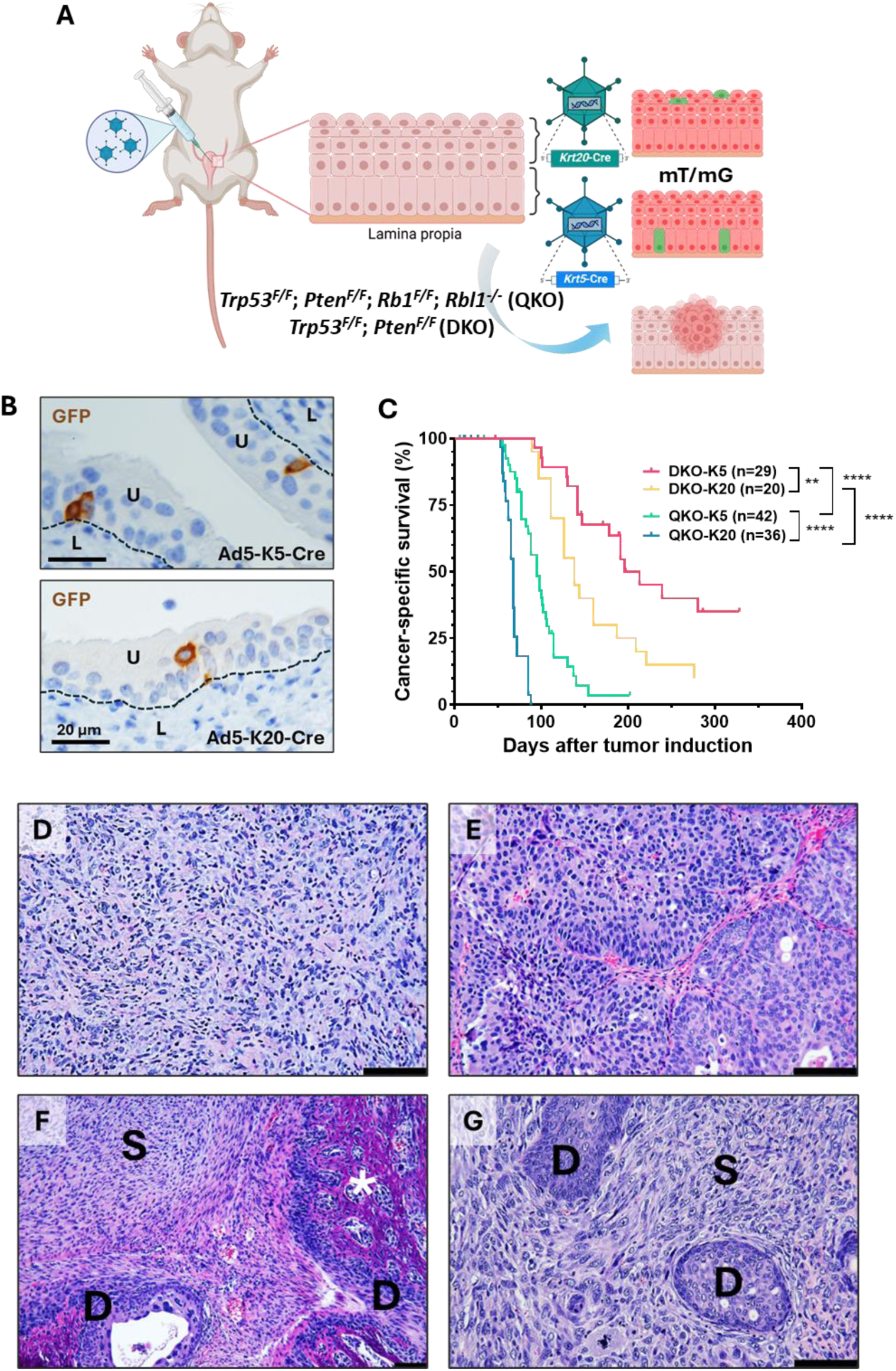
Development of invasive BC mouse models. **A.** Schematic representation of the generation of BC models. Adenoviruses (Ad5-Krt5-Cre and Ad5-Krt20-Cre) were injected into the bladder lumen of mTmG reporter mice and DKO and QKO mice for tumor induction. **B.** Representative images of GFP staining showing basal cells (Ad5-Krt5-Cre) and luminal cells (Ad5-Krt20-Cre) in the bladder urothelium of mTmG reporter mice. U = urothelium; L = lamina propia. Scale bar = 20 µm. **C.** Kaplan-Meier survival curve illustrating the percentage of cancer-specific survival following *Ad5-Krt5-Cre* and *Ad5-Krt20-Cre* injection in DKO and QKO mice. **p-value< 0.01; ****p-value< 0.0001. **D-G.** Representative H&E staining of the most common tumor morphologies observed in MIBC from the models: differentiated urothelial carcinoma (D), undifferentiated sarcomatoid carcinoma (E), and tumors with mixed patterns (F and G). Scale bar = 100 µm. Annotations: S = sarcomatoid pattern, D = differentiated pattern, * = bone metaplasia.

Cancer-specific survival ranged from 89 to 280 days following *Ad5-Krt5-Cre* or *Ad5-Krt20-Cre* administration in QKO and DKO mice. QKO mice had significantly shorter median survival than DKO mice for both K5-positive (95 vs. 213 days) and K20-positive cells (68 vs. 138 days) (p < 0.0001 in both cases). Additional differences were observed between DKO-K5 and DKO-K20 (213 vs. 138 days; p < 0.01), and between QKO-K5 and QKO-K20 groups (95 vs. 68 days; p < 0.0001) (**Figure 1C and Supplemental Table S1**). Tumor incidence ranged 80-90% in all groups, except for DKO-K5, which showed a lower incidence (55%), likely due to the prolonged latency period (**Supplemental Table S1**).

Histological analysis revealed that approximately half of the primary tumors in QKO mice (57.6% in QKO-K5 and 42.9% in QKO-K20) were high-grade, invasive undifferentiated carcinomas with malignant spindle cells, hyperchromatic nuclei, prominent nucleoli, high mitotic activity, and atypical mitotic figures, consistent with sarcomatoid carcinoma of the urinary bladder (20) (**Figure 1D, Supplemental Figure S2 and Supplemental Table S1**). The remaining tumors from QKO mice exhibited complete urothelial differentiation (around 10%) or a mixed pattern with both differentiated and undifferentiated regions (33.3% in QKO-K5 and 46.4% in QKO-K20) (**Figure 1, E–G, Supplemental Figure S2 and Supplemental Table S1**). In contrast, all tumors in DKO mice were invasive urothelial carcinomas with a uniform sarcomatoid morphology (**Figure 1D, Supplemental Figure S2 and Supplemental Table S1**).

We observed additional histological differences between groups, linked to Rb family alterations or the tumor cell origin. QKO tumors exhibited significantly higher levels of necrosis (41% vs. 2.9%; p < 0.0001), likely due to faster growth or reduced angiogenesis, along with increased immune cell infiltration (70.5% vs. 14.7%; p < 0.0001) and myxoid stroma presence (47.5% vs. 23.5%; p = 0.028), a feature commonly associated with sarcomatoid BC (21–23) (**Supplemental Figure S2, B–D, and Supplemental Table S2**). In contrast, pleomorphic giant cells, a characteristic associated with spindle-cell morphology in certain BC variants (24), were more frequently found in tumors derived from suprabasal cells (67.4% vs. 28.6%; p = 0.0002), with a trend toward higher frequency also observed in DKO tumors (**Supplemental Figure S2E and Supplemental Table S2**). Notably, unlike DKO tumors, a significant proportion of QKO tumors developed foci of osseous metaplasia with dystrophic calcification, particularly in basal-cell-derived tumors (57.6% vs. 35.7%; p = 0.0464), suggesting that Rb family loss promotes this abnormal differentiation, especially in cells with greater plasticity (**Supplemental Figure S2F and Supplemental Table S2**).

We also evaluated metastasis incidence across mouse models. Tumor cells primarily spread through the peritoneal cavity, leading to carcinomatosis (**Figure 2, A and B**), with the liver (**Figure 2C**), diaphragm (**Figure 2D**), spleen and pancreas (**Figure 2E**) as the most frequently affected organs. However, metastasis was also observed in distant organs, including the lungs, liver, and kidneys, through blood and/or lymphatic vessels (**Figure 2, F–H**). Notably, in both DKO and QKO mice, those inoculated with *Ad5-Krt5-Cre* exhibited a higher frequency tumor dissemination compared to those inoculated with *Ad5-Krt20-Cre* (62.5% vs. 16.7% in DKO, and 78.8% vs. 60.7% in QKO), with statistical significance observed only in the DKO group (p = 0.0122) (**Figure 2, A–B and Supplemental Table S1**). DKO-K20 tumors were the only ones that did not lead to distant metastasis, and no significant differences were observed in the other groups, with distant metastasis observed in 12.5% of DKO-K5, 30.3% of QKO-K5, and 21.4% of QKO-K20 (**Figure 2, A-B and Supplemental Table S1**). While no significant differences were observed in the proportion of tumor dissemination between the two models injected with *Ad5-Krt5-Cre*, mice in the QKO- K20 group exhibited a higher degree of dissemination compared to the DKO-K20 group (60.7% vs. 16.7%; p = 0.0055) (**Figure 2, A–B and Supplemental Table S1**). Furthermore, a trend toward a greater number of organs affected by carcinomatosis was observed in QKO mice compared to DKO mice (not shown). Except for one QKO-K5 mouse, which developed metastases with differentiated morphology in the liver (not shown), all other metastases, both carcinomatosis and distant, exhibited a sarcomatoid morphology, even when the primary tumor was fully differentiated. These metastases also exhibited several histological features similar to those of the primary tumors, such as myxoid stroma, abundant inflammation, necrosis, and bone metaplasia (**Figure 2, C–H**). Of note, the observed differences in survival are not explained by disparities in tumor dissemination. Likewise, differences in metastatic capacity cannot be ascribed to variations in the timing of tumor emergence across models.

**Figure 2.**
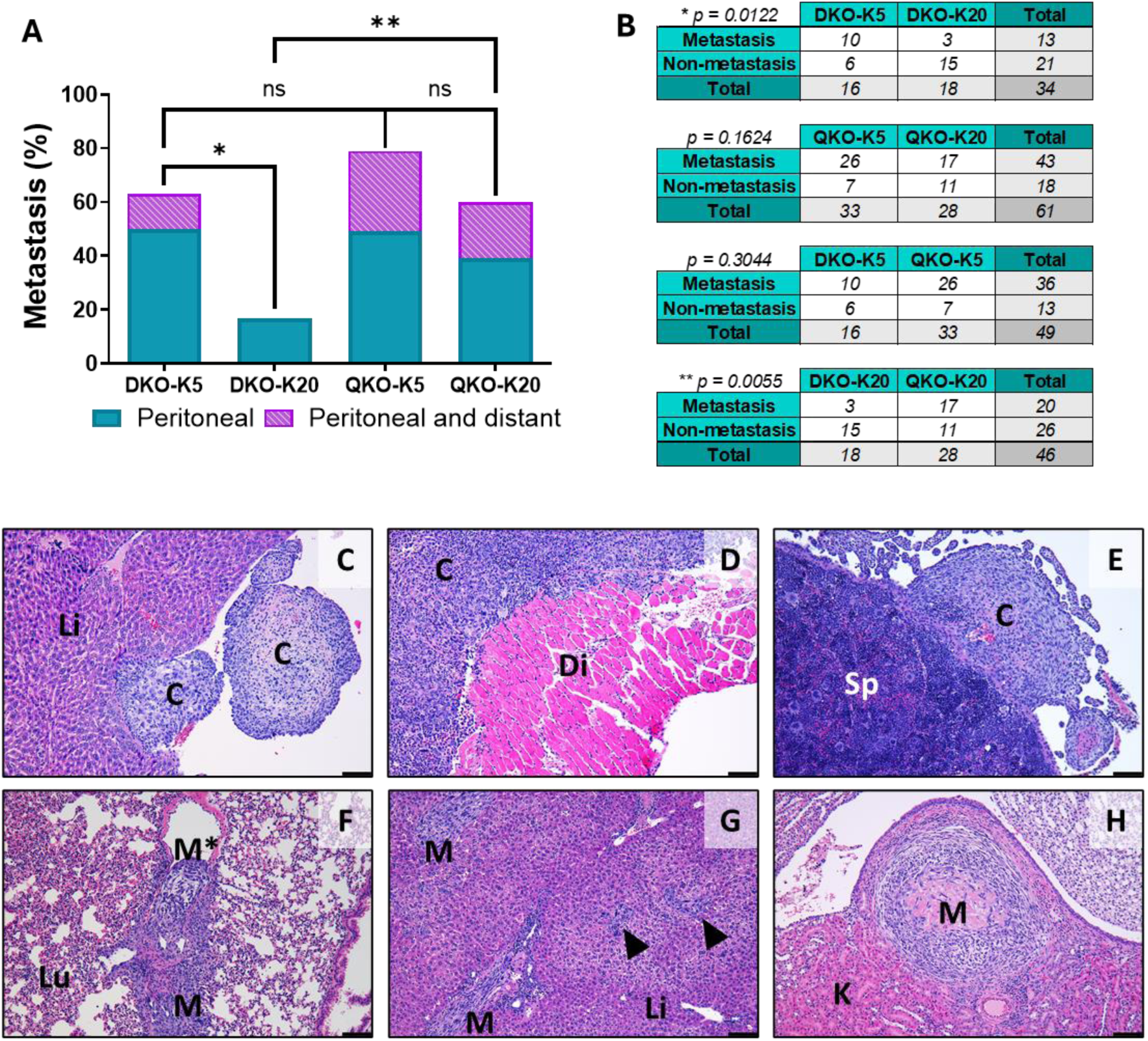
Metastasis analysis in invasive BC mouse models. **A.** Metastasis frequency (peritoneal and distant dissemination) across different mouse models. **B.** Contingency analysis of total metastases in mice with tumor development across groups. ns = not significant; *p-value< 0.05; **p-value< 0.01. **C-E.** Representative H&E staining of peritoneal metastases (carcinomatosis) in the most frequent locations such as the liver (C), diaphragm (D) and spleen (E). **F-H.** Representative H&E staining of distant metastases in frequent sites such as lung (F), liver (G) and kidney (H). Lu = Lung; Li = liver; K = kidney; M = metastasis; M* = metastasis within a vein. C = carcinomatosis. Di = diaphragm; Sp = spleen. Arrowheads indicate small tumor cell clusters. Scale bar = 100 µm.

Therefore, the different models generated recapitulate various morphological variants of human tumors, with distinct histological characteristics, differing onset times or growth rates, and varying incidences of dissemination.

### Transcriptomic analysis and molecular characterization of BC mouse models

To identify molecular pathways involved in tumor development and progression, as well as those underlying differences in tumor latency, morphology, and dissemination capacity across the different models, we performed a whole transcriptomic analysis. We analyzed five independent tumor samples from each mouse model, along with five healthy urothelial cell samples from both DKO and QKO mice (**Supplemental Table S3**). First, we compared the gene expression profiles of all tumor samples (n = 20) with those of healthy urothelial cells (n = 10). Principal component analysis (PCA) revealed two main expression profile clusters: one comprising all tumor samples and the other consisting of healthy urothelial cells, with no clear segregation according to the mouse model (**Figure 3A**). This comparison identified 8,478 significantly deregulated transcripts (fold change ±2, FDR p-value < 0.05), of which 5,163 were downregulated and 3,315 were upregulated in tumor samples (**Figure 3B and Supplemental Figure S3A**).

**Figure 3.**
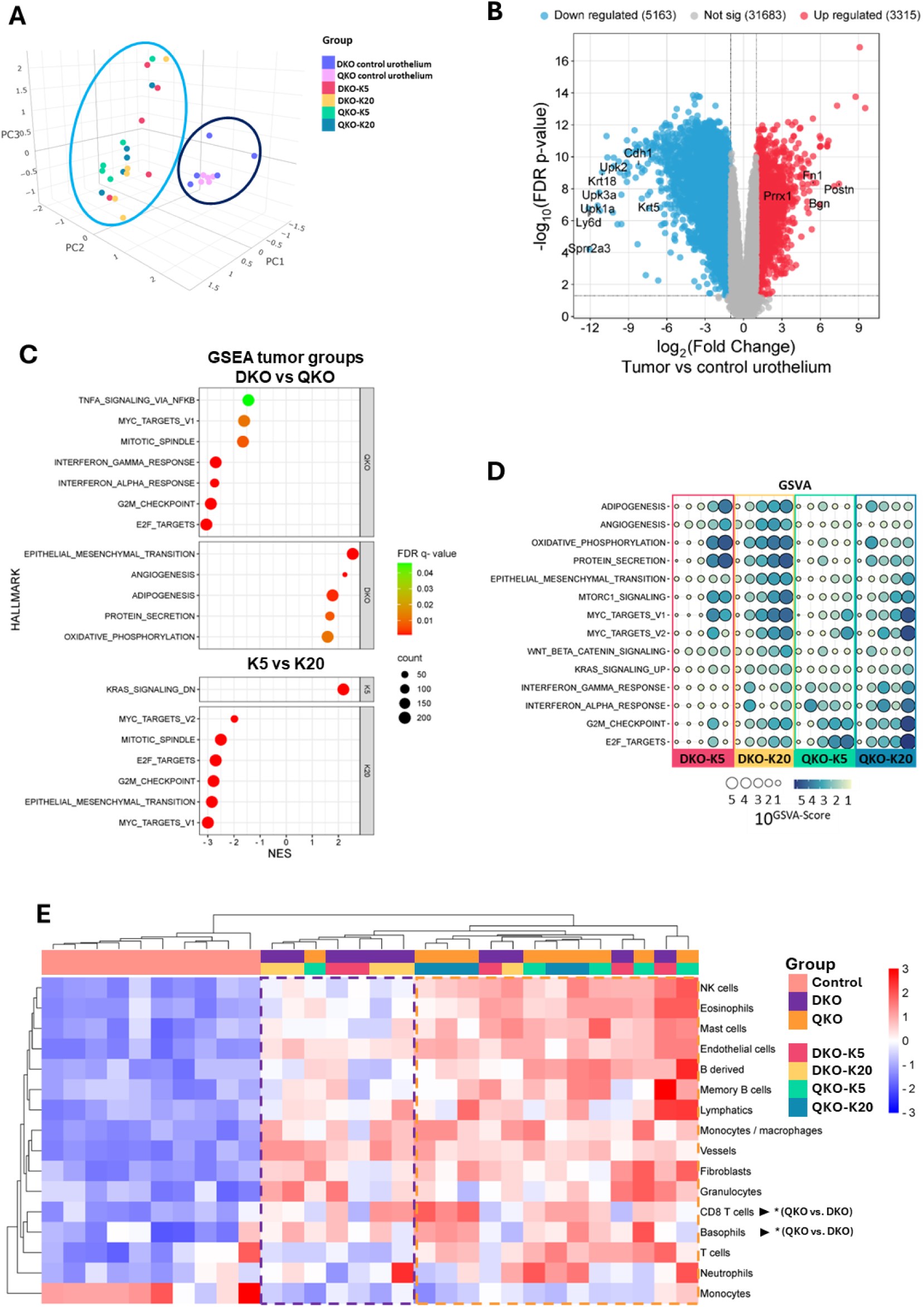
Transcriptomic analysis of tumors from different mouse models. **A.** Principal component analysis (PCA) showing the distribution of control urothelium and tumors from the different models. **B.** Volcano plot showing differentially expressed genes (fold change ± 2 and FDR p-value < 0.05) between control urothelium and tumors. **C.** GSEA identifying the most significantly enriched gene sets from the cancer hallmark molecular signature. Comparisons include QKO versus DKO tumors (top panels) and tumors derived from K5 versus K20 cells (bottom panels). Normalized enrichment score (NES), FDR q-value, and gene set size (count) are provided. **D.** GSVA scores for tumors across models, indicating key enriched gene sets from the cancer hallmark molecular signature. Scores are displayed as 10^GSVA-score^. **E.** TME analysis using mMCP-counter based on transcriptomic data, showing immune and stromal cell populations in tumors from each model.

Gene Set Enrichment Analysis (GSEA) comparing tumors and healthy urothelium revealed that tumors were enriched in gene signatures associated with epithelial-mesenchymal transition (EMT), cell cycle regulation (including G2/M checkpoint genes and E2F targets), angiogenesis, and immune microenvironment modulation, particularly inflammatory response, interferon-gamma response, and IL6/JAK/STAT3 signaling (**Supplemental Figure S3B**). Conversely, tumors exhibited decreased expression of gene signatures related to DNA repair, fat metabolism (including cholesterol homeostasis, adipogenesis, and fatty acid and bile acid metabolism), xenobiotic metabolism, and oxidative phosphorylation (**Supplemental Figure S3B**).

Interestingly, despite the behavioral and morphological differences among the tumor models evaluated, we did not observe clustering differences among them, nor between target cells or deleted genes (**Figure 3A and Supplemental Figure S3A**). Furthermore, differential expression analysis revealed no significantly deregulated transcripts (fold change ±2, FDR p-value < 0.05) across the models. Even when applying a less stringent threshold (non-adjusted p-value < 0.05), only a few significantly deregulated transcripts were identified, with no overlap among the different comparisons (**Supplemental Figure S3C**). Given these findings, we investigated whether the tumor models exhibited differences in the regulation of key biological pathways or processes related to tumor development and progression, despite the absence of statistically significant differentially expressed transcripts. These differences could potentially be driven by distinct transcriptional programs.

To address this, we performed GSEA comparing QKO vs. DKO tumors and tumors derived from K5-positive cells vs. those originating from K20-positive cells (**Figure 3C**). Additionally, we conducted Gene Set Variation Analysis (GSVA), an unsupervised method used to estimate variations in pathway activity at the individual sample level (**Figure 3D and Supplemental Figure S4**). Collectively, both analyses revealed that QKO tumors exhibited an enrichment in proliferation and cell cycle progression, consistent with their growth rates and histological features including increased mitotic spindle assembly, as well as a more active immune microenvironment, characterized by an enhanced interferon-alpha and interferon-gamma response (**Figure 3, C and D**). In contrast, tumors derived from DKO mice were enriched in EMT, angiogenesis, adipogenesis, and protein secretion pathways (**Figure 3, C and D**). Furthermore, tumors arising from K20-positive cells displayed increased expression of genes regulated by MYC, along with enhanced proliferation and EMT signatures (**Figure 3, C and D**). On the other hand, tumors originating from basal cells did not show enrichment in any relevant hallmark cancer gene sets (**Figure 3, C and D**). When analyzing specific subgroups, DKO-K20 tumors exhibited the strongest enrichment in EMT processes, whereas DKO-K5 tumors showed the lowest levels of proliferation- and immune response-related signatures (**Figure 3D**).

Given the observed differences in the proportion of tumors with abundant immune cell infiltration and variations in immune response profiles, we sought to further characterize the immune and stromal composition of the tumors using bulk transcriptomic data. We applied the mouse Microenvironment Cell Populations (mMCP) counter and performed an unsupervised clustering analysis, which revealed a clear distinction between urothelial controls and tumors (**Figure 3E**). Within the tumor samples, we identified two well-defined clusters: one with lower immune and stromal components, predominantly composed of tumors derived from DKO mice, and another with a higher immune cell presence, enriched in QKO tumors (**Figure 3E**). Compared to control tissue, all immune and stromal populations were significantly enriched in both tumor types, with the exception of neutrophils and T cells, which showed no differences between DKO tumors and controls (not shown). When comparing DKO and QKO tumors, most immune populations displayed higher abundance in QKO tumors, but only CD8^+^ T cells and basophils showed statistically significant increases (**Figure 3E**).

We further assessed the similarity of our models to established molecular subtype classification of advanced human bladder tumors, including the consensus classification, the Lund University algorithm and The Cancer Genome Atlas (TCGA). The consensus classification identified basal/squamous subtype as the most prevalent across most groups, while the stroma-rich subtype was predominant in the DKO-K20 group (**Figure 4A**). However, the Lund algorithm aligned better with tumor morphology, classifying differentiated tumors as basal/squamous-like and sarcomatoid tumors as mesenchymal-like (**Figure 4A**). Hierarchical clustering based on BC luminal and basal signature markers (25) showed that most undifferentiated or mesenchymal-like tumors clustered together, with subsets displaying either a double-negative profile or low basal marker expression. Differentiated tumors, however, clustered within a distinct subgroup characterized by basal marker expression and luminal marker downregulation (**Supplemental Figure S5A**). Given the high percentage of double-negative tumors, we evaluated sarcomatoid gene and microRNA expression signatures (26), which revealed a strong alignment with sarcomatoid profiles, distinguishing these tumors from control tissue (**Supplemental Figure S5, B and C**).

**Figure 4.**
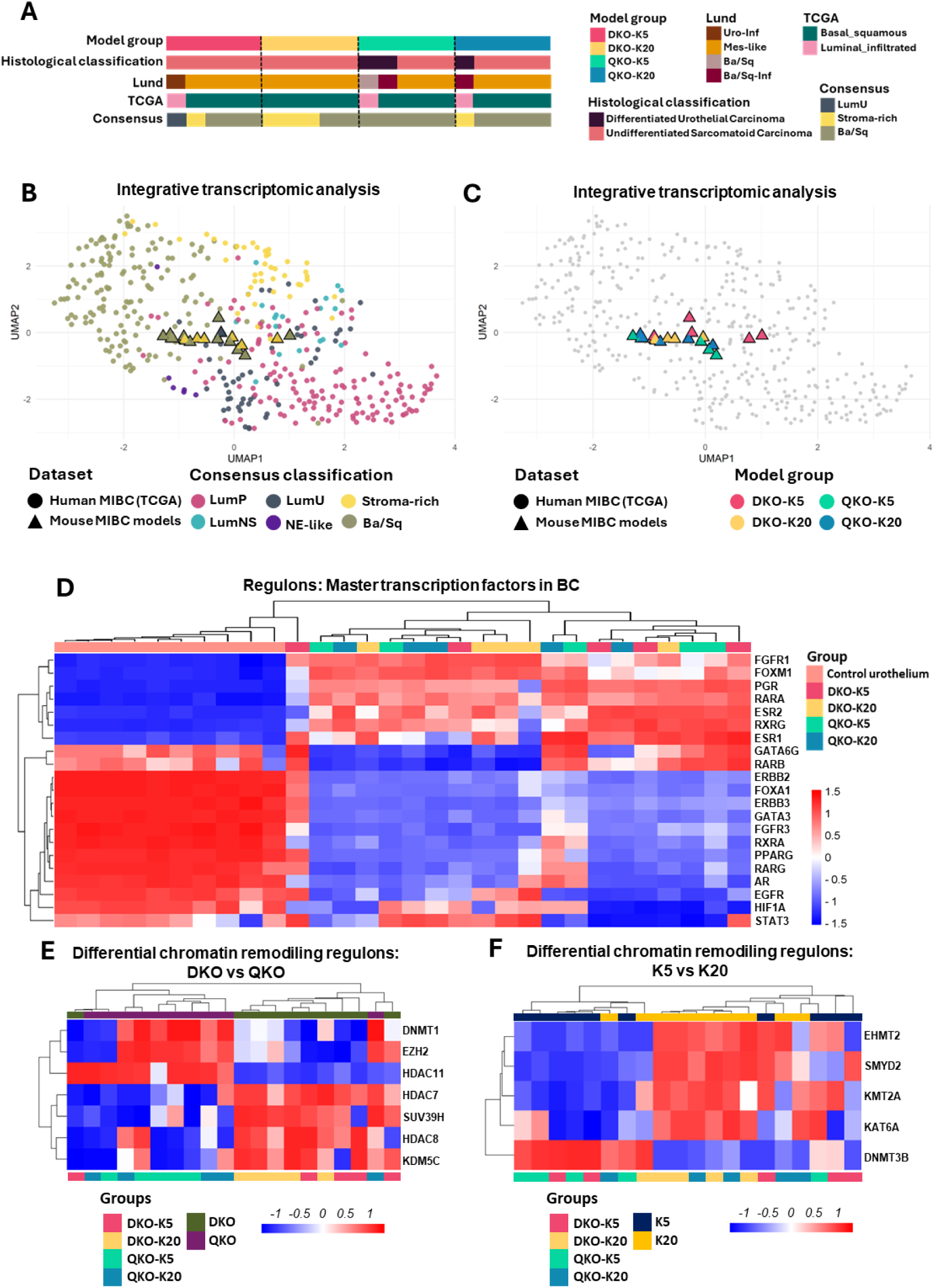
Molecular integrative clustering and regulon analysis. **A.** Molecular classification of tumors from each mouse model using human MIBC molecular classifiers. **B-C.** UMAP plot illustrating integrative transcriptomic analysis of tumors from different BC mouse models and MIBC samples from TCGA database. (B) shows tumor distribution according to the consensus molecular classification, while (C) displays distribution based on the originating mouse model. **D.** Heatmaps displaying hierarchical clustering of tumors across mouse models based on regulon activity of master transcription factors associated with BC. **E-F.** Differential regulon activity related to chromatin remodeling in (E) DKO vs. QKO tumors and (F) tumors derived from K5- vs. K20-positive cells.

An integrative transcriptomic analysis was performed to compare tumors from the mouse models with transcriptomic data from MIBC samples in the TCGA BLCA cohort (7). This analysis revealed that mouse tumors clustered within human tumors according to established molecular subtypes, independently of their model of origin (**Figure 4, Band C, and Supplemental Figure S6A**). Within this integrative framework, we further analyzed the distribution of mouse tumors using hierarchical clustering based on the top-500 most variable genes (**Supplemental Figure 6B**), luminal and basal BC signature markers (**Supplemental Figure S6C**), and the BC sarcomatoid gene expression signature (**Supplemental Figure S6D**). In all cases, we observed a primary segregation between human tumors classified as luminal (luminal papillary and luminal unstable) and those classified as basal/squamous, according to consensus classification. In the case of gene signatures, this segregation aligned with the expression of luminal versus basal markers, as well as with the upregulation or downregulation of sarcomatoid-associated genes (**Supplemental Figure S6, C and D**). Notably, mouse bladder tumors were distributed across the spectrum of human tumors, without clustering by model, reflecting a degree of molecular heterogeneity that parallels the diversity observed in human MIBC.

Consistently, we observed low activity of luminal-associated regulons, including FGFR3, FOXA1, GATA3, ERBB2/3, PPARG, RARG, RXRA, and AR (6,27). In contrast, most tumors exhibited increased activity of FOXM1, a regulon typically associated with the basal-like subtype (6,28), as well as RARA and PGR, which are linked to the stroma-rich subtype (6), and FGFR1 and ESR1, whose high activity has been associated with both molecular subtypes (6,29). Additionally, some tumors showed activation of other basal regulons, such as EGFR, STAT3, and HIF1A, or the stroma-rich regulon RARB (6) (**Figure 4D**).

Given the critical role of chromatin remodelling gene alterations in BC development and progression, we analyzed their regulatory activity across tumor models. Our analysis revealed widespread dysregulation of chromatin remodelers in tumors, with most displaying reduced activity compared to control tissue (**Supplemental Figure S7**). In tumor samples, we identified differential activity of seven remodelers between DKO and QKO tumors, all primarily involved in gene repression (**Figure 4E**), with the most pronounced differences observed in HDAC11, DNMT1, and EZH2 activity. HDAC11 activity was higher in QKO tumors than in DKO tumors (**Figure 4E and Supplemental Figure S7**), a pattern previously associated with BC progression and IL-10 repression (30,31), potentially contributing to the more inflammatory microenvironment observed in QKO tumors. Similarly, DNMT1 and EZH2 activities were also elevated in QKO tumors, showing a strong correlation. A direct interplay between these two methyltransferases has been demonstrated, linking them to tumor growth and migration (32,33), which aligns with the increased proliferation and metastasis observed in QKO models (**Figure 4E**). When assessing the influence of the cell of origin, the association was less pronounced, with differential activity detected in four methyltransferases (EHMT2, SMYD2, KMT2A, and DNMT3B) and one acetyltransferase (KAT6A) (**Figure 4F**). Among these, EHMT2 (also known as G9a) stood out due to its critical role in the regulation of cancer cell growth, particularly in BC (14), with higher activity observed in tumors originating from K20-positive cells.

Our analysis revealed transcriptional and epigenetic alterations across BC mouse models, highlighting key differences in proliferation, immune activity, and tumor plasticity.

### Establishment of syngeneic graft models of BC as preclinical tools for testing new therapeutic strategies

Despite the advantages of the developed GEMMs, their limitations, including unpredictable tumor onset, prolonged latency periods, and challenges in early tumor detection, led us to develop immunocompetent heterotopic models that allow for more precise tumor monitoring and synchronized growth patterns, enhancing the robustness of preclinical studies. To generate these models, we established primary cell cultures from bladder tumors in the four GEMMs (**Supplemental Figure S8, A and B**). These cells were characterized, confirming the deletion of the corresponding genes by genomic PCR and gene expression (**Figure 5, A and B, and Supplemental Table S3**). Furthermore, all cell lines exhibited a loss of epithelial and luminal markers while expressing vimentin, a mesenchymal marker, consistent with the expression profile observed in most primary tumors (**Supplemental Figure S9**).

**Figure 5.**
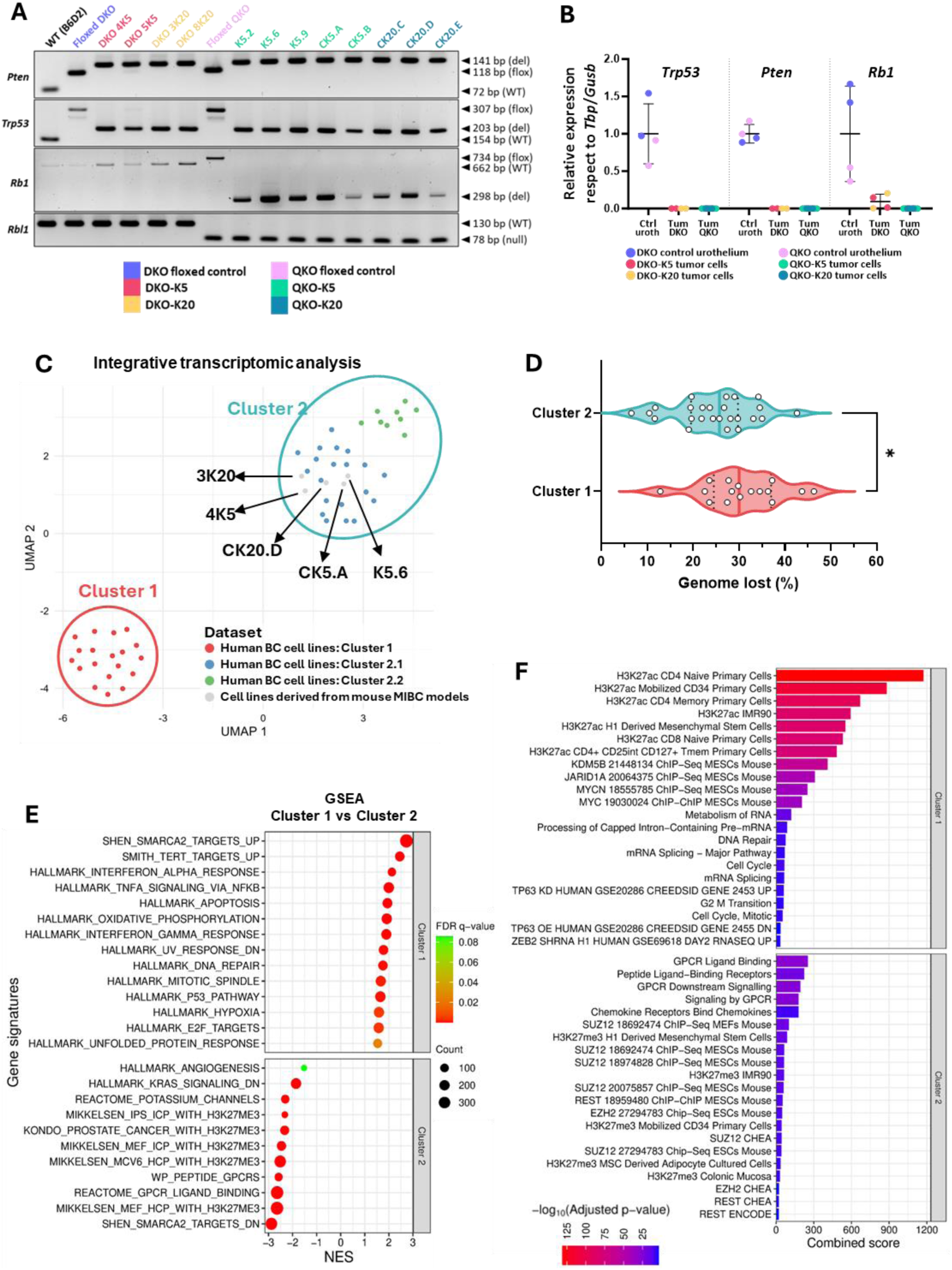
Characterization of tumor cell lines derived from different BC mouse models. **A.** PCR genotyping of the established cell lines for *Pten*, *Trp53*, *Rb1* and *Rbl1* genes. Genomic DNA from a wild type (WT) B6D2 mouse, as well as from DKO and QKO mice, served as control (see Supplemental Table 6 for details). WT = wild-type; floxed = allele carrying loxP site(s); deleted = allele resulting from loxP recombination; null = knockout allele containing a neomycin resistance cassette; bp = base pairs. **B.** Relative expression levels of *Trp53*, *Pten* and *Rb1* genes in the established cell lines. Each dot represents an individual cell line; data are presented as mean ± SD. **C.** UMAP plot displaying integrative gene expression-based clustering of mouse BC-derived cell lines and human BC cell lines. Names of the mouse BC cell lines are indicated. **D.** Comparison of genome loss percentages between the two major clusters of human BC cell lines. Individual values are shown; medians and interquartile ranges are indicated. *p-value< 0.05. **E.** GSEA identifying the most significantly enriched gene signature sets differentiating cluster 1 from cluster 2 in human BC cell lines. Normalized enrichment score (NES), FDR q-value, and gene set size (count) are provided. **F.** Bar plot showing enriched signatures identified using the Enrichr tool from differentially expressed genes between cluster 1 and cluster 2 of human BC cell lines. Combined score and adjusted p-value (–log₁₀) are displayed.

To assess the potential of these mouse cell lines and their resemblance to human BC models, we performed transcriptomic profiling of five mouse tumor-derived lines: one DKO-K5 (4K5) cell line, one DKO-K20 cell line (3K20), two QKO-K5 cell lines (K5.6 and CK5.A), and one QKO-K20 cell line (CK20.D) (**Supplemental Table S3**). These transcriptomic data were integrated with previously published transcriptomic data from 48 human BC cell lines (34), and a clustering analysis revealed two major transcriptional groups (**Figure 5C**), referred to as Clusters 1 and 2. Notably, Cluster 2, which includes all mouse BC cell lines, could be further subdivided into two subclusters, 2.1 and 2.2 (**Figure 5C**). Furthermore, we assessed the distribution of our mouse cell lines alongside human BC cell lines using hierarchical clustering based on the top-500 most variable genes (**Supplemental Figure S10A**), luminal and basal BC signature markers (**Supplemental Figure S10B**), and the BC sarcomatoid gene expression signature (**Supplemental Figure S10C**). Mouse cell lines were broadly distributed across the transcriptional landscape of human BC cell lines, clustering predominantly according to their genetic background (DKO or QKO) (**Supplementary Figure S10A–C**).

Analysis of known genomic alterations in these human cell lines revealed that Cluster 1 harbored a significantly higher proportion of genomic deletions compared to Cluster 2 (p < 0.05; **Figure 5D**). Furthermore, differential expression analysis followed by GSEA and Enrichr enrichment revealed that Cluster 1 was significantly enriched for genes involved in interferon-alpha and -gamma response pathways, TNF-alpha signaling, cell cycle progression, mitosis, and proliferative programs. This cluster also displayed loss of TP63 expression, DNA repair defects, activation of TERT target genes, and strong upregulation of MYC and RNA processing and splicing machinery (**Figure 5, E and F**). Strikingly, Cluster 1 showed robust enrichment for chromatin remodeling signatures, including activation of the histone demethylases JARID1A and KDM5B (JARID1B), as well as the SWI/SNF complex, particularly through the ATPase subunit SMARCA2, known to associate with open chromatin (H3K27ac) and active transcription. In parallel, there was strong evidence of Polycomb repressive complex 2 (PRC2) activity, marked by activation of EZH2 and SUZ12 and repression of H3K27me3 target genes (**Figure 5, E and F**). While SWI/SNF and PRC2 are often functionally antagonistic at the same genomic loci, certain cellular context may exhibit coexistence of chromatin-opening and -closing signals, reflecting transcriptional plasticity, particularly in cancer (35,36). In contrast, Cluster 2 showed low activity of chromatin remodeling complexes and was instead characterized by strong activation of G-protein coupled receptor (GPCR) signaling, particularly through ligand and peptide binding, as well as chemokine receptor activation (**Figure 5, E and F**). Interestingly, within Cluster 2, subcluster 2.1, which harbors increased similarities with the mouse tumor-derived cell lines, displayed a significantly higher proportion of *TP53*-mutant cell lines compared to subcluster 2.2 (p < 0.05; not shown).

Next, to assess the engraftment potential of these primary cell cultures, we performed subcutaneous injections of ten different tumor cell lines into syngeneic mice. All tested DKO cell lines achieved 100% engraftment, albeit with varying growth rates. In contrast, not all QKO- derived tumor cells fully engrafted and between 50% and 100% exhibited tumor regression, with the most pronounced effect observed in QKO-K20 cells (**Supplemental Figure S11A and Supplemental Table S3**). Based on their engraftment efficiency and appropriate growth dynamics for preclinical testing, we selected two cell lines derived from K5-driven tumors: 4K5 from a DKO tumor and K5.6 from a QKO tumor, for subsequent *in vivo* experiments.

Histological analysis of syngeneic graft tumors from these two cell lines confirmed that they retain the sarcomatoid features observed in primary tumors (**Supplemental Figure S11B**). Global transcriptomic analysis further revealed that these tumors exhibit a gene expression profile similar to the specific primary tumors from which were derived, including luminal, basal, and sarcomatoid gene signatures, supporting the conservation of primary tumor characteristics (**Supplemental Figure S11C**). However, no metastatic dissemination was detected in these models, most likely due to the aggressive and rapid tumor growth of these subcutaneous graft-based systems. Together, these findings demonstrate the successful development of transplantable murine MIBC cell lines capable of serial transplantation in immunocompetent syngeneic hosts and their potential use as preclinical tools.

In our previous research, we demonstrated that the antitumor activity of the CDK4/6 inhibitor Palbociclib in human BC was independent of Rb status (15). Moreover, considering the potential synergistic interaction between CDK4/6 inhibitors and ICIs (37,38), we further explored this combination in our experimental systems. We evaluated the cytotoxic effects of palbociclib in selected primary cell cultures derived from DKO (4K5) and QKO (K5.6) models. Both DKO and QKO tumor cells exhibited comparable sensitivity to palbociclib, with IC_50_ values falling within a narrow range (4K5 = 2.81 μM; K5.6 = 1.93 μM) (**Figure 6A**). Furthermore, both cell lines showed a significant induction of apoptosis, as assessed by Annexin V/DAPI staining (**Supplemental Figure S12A**). We next examined the impact of palbociclib on cell cycle progression and its association with Rb status. Treatment with palbociclib in Rb wild-type cells led to a significant reduction in S-phase entry, indicating G1 phase arrest (p < 0.01). In contrast, Rb-deficient cells displayed an increase in the G2/M phase (p < 0.05), accompanied by a significant reduction in S phase (p < 0.05) (**Figure 6B**). Additionally, we evaluated PD-L1 expression following palbociclib treatment in both cell lines and observed a marked increase in both the percentage and intensity of PD-L1-positive live cells (**Figure 6C**). Given these observations, we proceeded to evaluate palbociclib treatment either alone or in combination with a blocking anti-PD-L1 antibody (avelumab). We first conducted a pilot experiment in a small cohort of DKO mice injected with Ad5-K5-Cre (5 mice per group, including a control group), in which the combinatory treatment resulted in evident tumor cell death and increased immune cell infiltration compared to palbociclib monotherapy (**Supplemental Figure S12B**).

**Figure 6.**
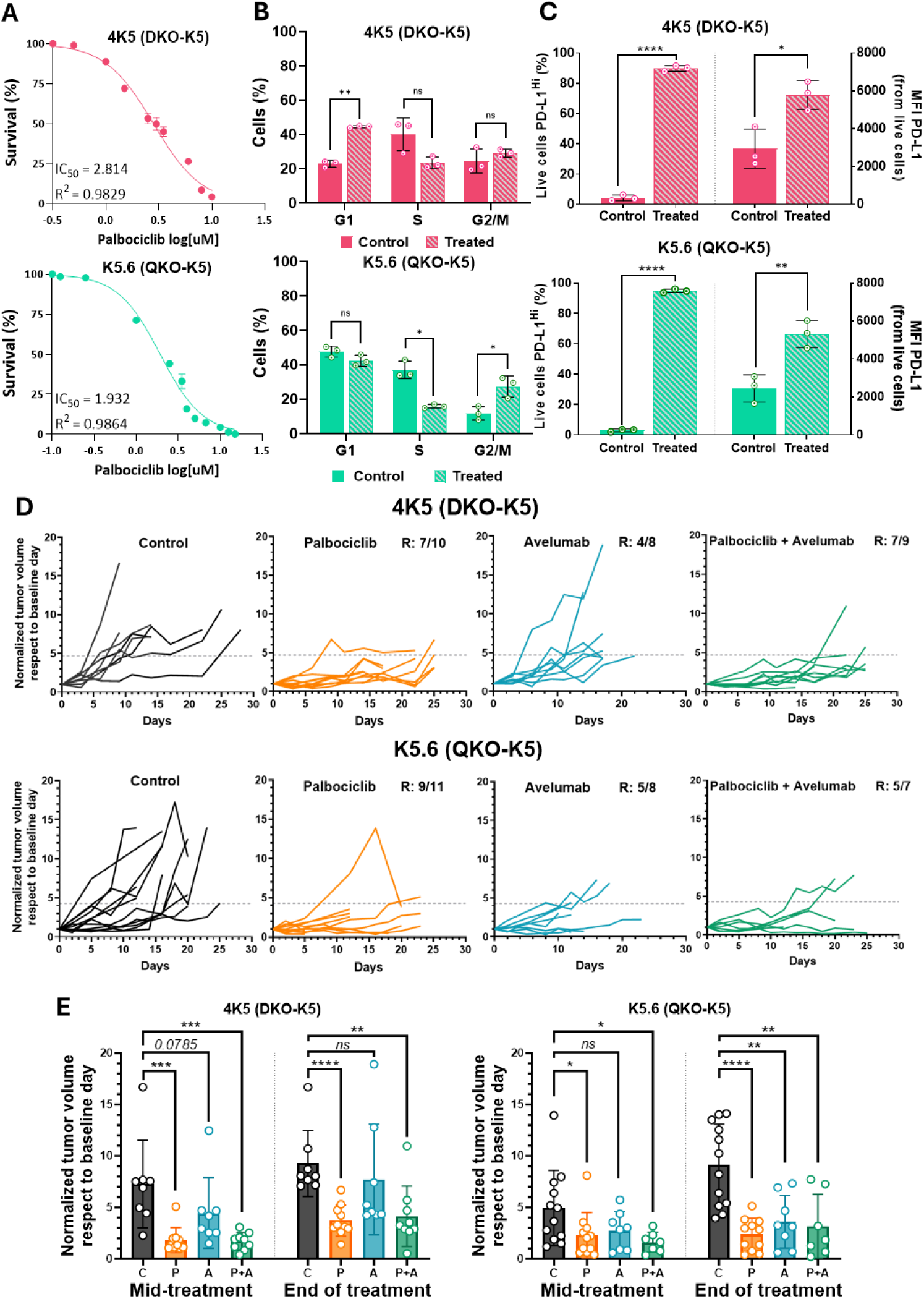
Immunocompetent syngeneic graft models of BC as preclinical tools for evaluating palbociclib combined with avelumab. A-C. *In vitro* analysis of mouse BC cell lines derived from K5-positive cells isolated from DKO (4K5; top) and QKO (K5.6; bottom) tumors: (A) sensitivity to palbociclib assessed by XTT assays. Data are presented as mean ± SEM; (B) Cell-cycle changes following palbociclib treatment for 48 hours at the IC_50_ analyzed by flow cytometry; (C) Induction of PD-L1 expression after palbociclib treatment, depicting the percentage of live cells with high PD-L1 expression (left axis) and the mean fluorescence intensity (MFI) of PD-L1 (right axis), as assessed by flow cytometry. In B and C, data are presented as mean ± SD from three independent experiments, with the average value of each experiment shown as a single data point. **D.** *In vivo* evaluation of palbociclib monotherapy or combined with avelumab (anti-PD-L1) using immunocompetent syngeneic graft models of BC (DKO-K5, top; QKO-K5, bottom). Spider plots show normalized tumor growth curves for each treatment group relative to baseline. The dotted line indicates the threshold for non-responders, defined as half the average tumor volume of the control group. R = responders. **E.** Comparison of normalized tumor volumes (relative to baseline) between treatment groups at mid-treatment and the end of treatment for both models. Data from each individual are shown, along with the mean ± SD. C = control; P = palbociclib; A = avelumab; P+A = combination palbociclib plus avelumab. ns = not significant; *p-value< 0.05; **p- value< 0.01; ***p-value< 0.001; ****p-value< 0.0001.

However, as previously mentioned, challenges related to follow-up and the broad time window for tumor development hindered the consistency of these experiments. To address these limitations, we turned to the syngeneic mouse models established with K5.6 and 4K5 cells, as previously described. In these models, we followed the treatment schedule outlined in **Supplemental Figure S13A**, ensuring the best possible synchronization of tumor growth and treatment administration. Tumor growth analyses showed that both mouse models responded well to palbociclib, alone or with a PD-L1 inhibitor, while PD-L1 monotherapy was less effective (**Figure 6D**). In the DKO models, palbociclib (± avelumab) was consistently effective, whereas avelumab alone showed only a transient, non-significant effect (**Figure 6E**). In QKO tumors, responses to palbociclib (± avelumab) strengthened over time, and avelumab alone became effective only at the end of treatment (**Figure 6E**).

Furthermore, analysis of the TME in DKO-K5-derived tumors after treatment revealed a notable increase in CD45⁺ immune cell populations in both groups treated with palbociclib. This was accompanied by a pronounced decrease in the CD45⁻GFP⁺ population, corresponding to tumor cells, indicating effective tumor cell clearance. Focusing on the lymphoid compartment, we observed a reduction in B cells, CD8⁺ and CD4⁺ T lymphocytes, and NK cells, with statistically significant decreases observed only in CD8⁺ T cells and NK cells. Interestingly, all treatment groups exhibited an expansion of the myeloid lineage (defined in our panel as lineage-negative after exclusion of other microenvironment populations), which was GFP⁺, suggesting a high level of phagocytosis of tumor cell debris. This increase was statistically significant in the groups treated with palbociclib exclusively. Moreover, palbociclib also affected stromal cell populations, leading to a reduction in fibroblasts within the tumors, specifically decreasing the proportion of myofibroblastic cancer-associated fibroblasts (myoCAFs), as well as CD31⁺ endothelial cells (**Supplemental Figure S13B**).

These findings highlight the utility of our cell lines and *in vivo* models for testing therapies in immunocompetent settings with distinct TME profiles.

## DISCUSION

BC is a prevalent malignancy for which impactful biological discoveries and therapeutic advances have lagged behind those seen in other cancer types. Although the advent of immunotherapies has led to improved survival in some patients, the proportion of individuals who derive objective clinical benefit remains limited. Moreover, treatment options for metastatic disease are still scarce. Preclinical models of BC continue to be essential for uncovering novel biological mechanisms and developing new therapeutic strategies. While *in vitro* systems offer a controlled environment to study therapeutic responses, they lack a representative TME, thus reducing the impact of therapies in particular those affecting the dynamic immune system. *In vivo* models, such as BBN-induced bladder tumor model, provide a more physiologically relevant context; however, tumor incidence and phenotype are highly dependent on the genetic background of the mouse strain used (13). Furthermore, GEMMs of urothelial carcinoma, especially those that recapitulate muscle-invasive and metastatic disease, remain underrepresented compared to GEMMs of other cancer types. A significant limitation in developing bladder-specific GEMMs lies in the paucity of genes uniquely expressed in the urothelium, which hampers targeted gene recombination in relevant bladder cell populations. This challenge has been partially addressed through the intravesical delivery of Cre-expressing adenoviruses, allowing the use of promoters such as keratin K5 and K20, which, while not bladder-specific, enable targeted recombination in basal and non-basal urothelial cells. This approach facilitates direct comparisons of tumor initiation across distinct urothelial lineages and allows for efficient gene targeting in both male and female mice, circumventing the technical limitations associated with transurethral delivery in males.

We focused our studies on the inactivation of the tumor suppressor genes *Pten* and *Trp53*, which has previously been shown to be sufficient to drive BC tumorigenesis. However, our findings diverge in some aspects from prior reports. In line with previous data, we demonstrated that concurrent elimination of the Rb family, specifically to prevent functional compensation of *Rb1* loss by *Rbl1*, promotes a more basal-like differentiation program that rapidly progresses to a sarcomatoid phenotype. In contrast, *Pten*/*Trp53* inactivation alone gave rise almost exclusively to sarcomatoid tumors. Notably, different groups using the DKO model with CMV-Cre adenoviruses have reported the development of both differentiated tumors (39) and sarcomatoid tumors resembling those observed in our study (40). While the underlying reasons remain unclear, strain-specific factors and context-dependent somatic events triggered upon inactivation of these tumor suppressor genes may contribute to the observed differences. Importantly, our promoter-specific recombination studies in basal and suprabasal urothelial cells using K5 and K20 promoters revealed highly similar phenotypes, strongly suggesting that ectopic Cre expression is not responsible for the divergence. Additionally, the discrepancies in tumor incidence and aggressiveness between our models and those reported using Ck5- CreERT2-driven recombination in DKO urothelial cells may reflect differences in the recombination strategy. As previously discussed, bladder tumors induced via transgenic Cre alleles tend to exhibit less aggressive behavior but follow a more predictable developmental course (41).

Our results clearly demonstrate that tumor phenotypes, including incidence, histopathological features, and transcriptional alterations, are profoundly influenced not only by the specific tumor suppressor genes inactivated, but also by whether recombination is directed to basal or suprabasal urothelial cells. However, none of these genetic combinations led to the development of luminal tumors, suggesting that additional genetic drivers are required for the emergence of these BC subtypes in mice. This is consistent with the molecular characterization of human BC, which implicates other pathways in the pathogenesis of luminal disease. Supporting this notion, we observed minimal activation of canonical luminal regulons such as FGFR3, FOXA1, GATA3, and PPARG. Strikingly, our models also revealed notable differences in metastatic behavior. While all models exhibited peritoneal dissemination, likely via carcinomatosis, distant metastases to the lung and/or liver were more frequent in tumors arising from recombination in K5-expressing cells. This suggests that basal cells may have a greater intrinsic potential for progression toward visceral dissemination. Nevertheless, our transcriptomic profiling of primary tumors, while revealing molecular differences across models, did not fully account for the divergent metastatic patterns observed. For instance, signatures of EMT were more pronounced in DKO-K20 tumors, which notably lacked distant metastases. This indicates that additional, currently unidentified factors likely contribute to the metastatic potential of basal-derived tumors.

Another notable difference among the models lies in their distinct inflammatory microenvironment. Transcriptomic analyses using GSVA and mMCP-counter deconvolution revealed that tumors from QKO models are enriched in immune-related signatures and exhibit a higher infiltration of immune cells, aligning with histopathological observations. The relationship between this immune-enriched microenvironment and the deletion of the Rb family, as well as its potential contribution to metastatic progression, remains to be elucidated and is currently under investigation.

The GEMMs developed in this study provide robust platforms to investigate the molecular bases of aggressive BC. However, variability in tumor penetrance and latency across models presents a significant limitation for the consistent evaluation of preclinical therapies, as previously observed when relying exclusively on QKO K5-derived primary tumors (14,15). To overcome these limitations, we established a series of mouse tumor cells lines and syngeneic graft models from both DKO and QKO tumors. Given the ongoing debate surrounding the influence of *Rb1* status in modulating sensitivity to CDK4/6 inhibitors (15), we directly compared the therapeutic response of DKO and QKO models. Our findings indicate that palbociclib induces marked growth arrest *in vitro* and *in vivo* across both models, independent of *Rb1* status. In light of recent reports suggesting synergy between CDK4/6 inhibitors and ICIs, we leveraged the immunocompetent nature of our syngeneic models to assess the efficacy of this combination in vivo. Anti–PD-L1 monotherapy demonstrated greater effectiveness in the QKO model, likely reflecting its previously reported higher baseline immune infiltration. However, we did not observe an additive or synergistic effect from the combination therapy, potentially due to the robust antitumor activity of palbociclib alone at the administered dose. These findings suggest that the response to anti–PD-L1 therapy may be influenced by the pre-existing immune landscape of the tumor, and that optimizing combination regimens—such as using lower doses of palbociclib—could help mitigate toxicity while preserving therapeutic benefit. Overall, the cell lines and *in vivo* models generated in this study represent valuable resources for evaluating therapeutic strategies and dissecting BC responses in immunocompetent settings with distinct TME profiles.

In summary, this study provides a comprehensive framework to investigate the complex biology of MIBC and mBC through the development of refined GEMMs, syngeneic models, and tumor-derived cell lines. By integrating genetically defined *in vivo* systems with immunocompetent settings, our work addresses critical limitations in the field, including the lack of experimental models that faithfully recapitulate the heterogeneity, immune landscape, and metastatic potential of advanced disease. These models enable direct interrogation of how specific genetic alterations and cellular contexts influence tumor behavior, therapeutic responses, and immune interactions. Moreover, they offer a unique platform to explore mechanisms of resistance, identify predictive biomarkers, and optimize combination therapies involving immunomodulatory agents. The tools and insights generated through this work will be instrumental in advancing both the biological understanding and translational development of effective treatments for aggressive urothelial carcinoma.

## METHODS

### Genetically engineered mouse models and Cre-expressing adenoviruses

Conditional Quadruple Knock-Out (QKO; *Pten^flox/flox^, Trp53^flox/flox^, Rb1^flox/flox^*, *Rbl1*^-/-^) and Double Knock-Out (DKO; *Pten^flox/flox^, Trp53^flox/flox^)* mice were generated in-house through conventional breeding of strains obtained from collaborating institutions (Netherlands Cancer Institute, Massachusetts Institute of Technology, Stanford University), resulting in an immunocompetent mixed FVB/129S genetic background. Adenoviruses expressing Cre recombinase under the control of keratin 5 or keratin 20 promoters were obtained from the Viral Vector Production Unit at the Universitat Autònoma de Barcelona and were surgically injected into the bladder lumen, as previously described (42,43). Tumor development was routinely monitored by manual palpation. The bovine keratin 5 promoter was characterized by our group (18,19) and has been extensively used in published mouse models. In contrast, the murine keratin 20 promoter was specifically cloned for this study, as detailed in the *Supplemental material*. Experiments were conducted in male and female mice aged 8–12 weeks.

### *In vivo* validation of K5 and K20 promoters

Adenoviruses encoding Cre recombinase under the control of K5 and K20 promoters were inoculated as previously described intoB6.129(Cg)- *Gt(ROSA)26Sortm4^(ACTB-tdTomato,-EGFP)Luo^*/J mice, obtained from the F.X. Real laboratory at CNIO (Spain). These mice harbor loxP sites flanking a membrane-targeted tdTomato (mT) cassette and exhibit strong red fluorescence in all examined tissues and cell types. Cre-mediated recombination excises the mT cassette, allowing expression of the downstream membrane-targeted EGFP (mG) cassette. Six days after injection of the Cre-expressing adenoviruses, mice were euthanized, and bladders were harvested and processed as described in the *Histological Analysis* section (*Supplemental material*). Entire bladders were sectioned at 3–5 µm thickness and stained with an anti-GFP antibody (**Supplemental Table S5**) following standard immunohistochemistry protocols.

### Primary cell cultures

For the establishment of primary cell cultures, tumor specimens obtained from the different GEMMs were washed and minced into small pieces. Tumor dissociation was carried out by enzymatic digestion with 200 µg/mL of collagenase P (11213857001 Roche), 800 µg/mL dispase II (D4693 Sigma-Aldrich), and 100 µg/mL DNase I (10104159001 Roche) at 37°C, mixing samples every 20 minutes until the sample was disaggregated. Disaggregated tissue was filtered through a 40 µm strainer and centrifuged at 16 g for 7 minutes. Cells were cultured with DMEM *GlutaMAX™* (Gibco-BRL Life Technologies) with 10% FBS (Gibco-BRL Life Technologies) and 1% antibiotic-antimycotic (Gibco-BRL Life Technologies) at 37°C in a humidified atmosphere of 5% CO2.

### Syngeneic graft mouse model

To generate syngeneic graft mouse models, 3x10^6^ cells in total volume of 100 μL of Dulbecco’s Modified Eagle Medium (DMEM) were injected subcutaneously into both flanks of 12- to 16-week-old mice. Four animals (two females and two males) were used for each cell line, employing the appropriate mouse strain for each case (DKO or QKO). The tumor growth was monitored 2-3 times a week using an electronic gauge. The volume of the tumors was calculated as 4π/3 x (length/2) x (width/2)^2^.

### Drugs and *in vivo* treatments

Palbociclib (CDK4/6 inhibitor) and avelumab (human anti-PD-L1 that recognized murine epitope) were provided by Pfizer SLU España (CPT Grant Request ID: 60314347, Project ID: WI235570). For *in vitro* studies, palbociclib was dissolved and stored in dimethyl sulfoxide (DMSO) at the concentration of 100 mM at -20°C. Intermediate solutions at 10 mM were prepared in PBS and stored at -20°C until further use. Working solutions in PBS were further prepared (100-1000 µM) at 1% of DMSO. The final solutions (1-10 µM) in the medium were all freshly prepared before treatments. The final concentration of DMSO was 0.01% and maintained constant. For *in vivo* studies, palbociclib was dissolved in PBS at a working concentration of 20 mg/mL, and avelumab was dissolved with PBS at a final concentration of 2 mg/mL. Mice treatment was started at the time of tumor reached 100-150 mm^3^ using a regime of 40 mg/kg of palbociclib injected intraperitoneally five days per week and/or avelumab once per week for a total of three injections of 200 µg per injection (**Supplemental Figure S13A**). Male and female animals aged 12 to 16 weeks were used for each cell line, employing the appropriate mouse strain in each case (DKO or QKO). Cages were randomized to experimental groups to minimize environmental bias. Animals were humanely euthanized upon reaching predefined humane endpoints, primarily due to tumor size or the development of tumor-associated ulceration.

### Gene expression microarray analysis

Gene expression microarray analysis was performed using 12 ng of total RNA from each fresh tissue sample (control urothelium) or 30 ng from FFPE tumor samples. Transcriptomic profiling was carried out with the Clariom D Assay Mouse (Affymetrix, Thermo Fisher Scientific) and the GeneChip WT Pico Kit (Thermo Fisher Scientific), following the manufacturer’s protocols. Double-stranded cDNA synthesis included 9 PCR cycles, selected based on the *in vitro* transcription cycling guidelines. Data analysis was conducted using Transcriptome Analysis Console (TAC) v4.0.2 (Thermo Fisher Scientific). Expression data were normalized, and both background and batch effects were corrected using Guanine Cytosine Count Normalization (GCCN) and Signal Space Transformation (SST). Data summarization was done using the Robust Multi-Array Average (RMA) algorithm, implemented in the TAC software. Probe annotation was based on the mm10 (*Mus musculus*) genome version. Differentially expressed genes were identified using the eBayes ANOVA method.

### RNA-seq library preparation and sequencing

RNA sequencing analysis was performed using total RNA extracted from mouse BC cell lines, processed into biological duplicates. Stranded mRNA libraries were prepared with the TruSeq Stranded mRNA kit (Illumina), following the manufacturer’s protocols. Pairwise sequencing (2 × 150 bp) was performed on a NovaSeq 6000 platform (Illumina) at the Centro Nacional de Análisis Genómico (CNAG, Barcelona, Spain), obtaining more than 60 million reads per sample. Adapter trimming and quality filtering of the FASTQ files were performed with Trimmomatic. The quality of the reads was evaluated with FastQC. Alignment to the GRCm38 mouse reference genome (mm10) was performed with STAR aligner. Quantification at the gene level was performed with featureCounts. Counts were normalized using DESeq2’s median-of-ratios method.

### Bioinformatics analysis

PCA was performed using the normalized gene expression matrix of BC mouse tumors obtained from TAC 4.0 software, employing the *prcomp* function in R (https://www.r-project.org/), and the results were visualized with the *plotly* package. Volcano plot displaying differentially expressed genes between control urothelium and tumors was generated using *SRplot* (44). GSEA was performed using GSEA software (v4.1.0) and the Molecular Signature Database (MSigDB) with the *Hallmarks* gene set collection (*h.all.v7.5.symbols.gmt*). The analysis was conducted with 1,000 permutations (45). Additionally, GSVA was performed using the *GSVA* package in R (v1.42.0) with the *Hallmark* gene set (46). Results from both GSEA and GSVA were visualized as bubble plots using *SRplot* (44).

Mouse tumor samples were classified according to two published human MIBC molecular classifications (Lund and TCGA) using the *BLCAsubtyping* package in R. Additionally, the consensus molecular classification of MIBC was assigned using the *consensusMIBC* package (6). Since these classification algorithms were developed for human transcriptomic data, human orthologs were mapped and used for the molecular subtyping of mouse tumor samples when discrepancies in gene nomenclature were present. To further characterize the TME, the *mMCPcounter* package in R was used to deconvolute transcriptome data from each sample, estimating cell type abundance scores correlated with cell-type proportions.

Heatmaps and hierarchical clustering analyses of differentially expressed genes, gene signatures, regulons, and mMCP-counter data were generated using *SRplot* (44). Hierarchical clustering was performed using the Euclidean distance metric and complete linkage method. For the comparison of gene signatures between mouse primary tumors and syngeneic graft tumors, heatmap was generated using Multi-Experiment Viewer (MeV 4.9.0). In this case, normalized gene expression values were log2-transformed, and expression ranges were adjusted on a per-gene basis to enhance visualization.

Venn diagram was generated to illustrate the overlap of differentially expressed genes across comparisons of the different BC mouse models (fold change ±2 and p-value < 0.05). The diagram was created using the online tool available at https://bioinformatics.psb.ugent.be/webtools/Venn/.

We performed integrative cross-species and cross-platform analyses by combining our transcriptomic data from mouse bladder tumors obtained by microarrays with RNA-seq data from human invasive tumors (TCGA BLCA cohort), as well as our RNA-seq data from mouse BC cell lines with previously published microarray data from human BC cell lines (34). Prior to integration, gene identifiers were harmonized using human-mouse orthologs. To mitigate batch effects and improve comparability, expression matrices were independently transformed into ranks per sample and z-scores were standardized across genes following a Rank-based INtegration (RANK-IN) approach (47) (http://www.badd-cao.net/rank-in/). UMAPs were performed on the combined matrices using the *uwot* R package to visualize sample clustering and transcriptomic similarity. Heatmaps were generated using the *pheatmap* package in R, showing the most variable genes or specific gene signatures. Rows were clustered by Euclidean distance metrics and columns by Pearson’s correlation, both using Ward’s D2 method.

Differential gene expression analysis between human BC cell line clusters was performed using the *limma* package in R (v4.3.1), based on the RMA-normalized expression matrix (34). Linear modeling was applied using a design without intercept (∼0 + group), and contrasts were computed between each cluster and the remaining samples. Moderated statistics were obtained using empirical Bayes shrinkage (eBayes). GSEA was performed using the GSEA software (v4.1.0) and the Molecular Signature Database (MSigDB) employing the *Hallmark* gene set collection (*h.all.v7.5.symbols.gmt*) and the *Chemical and Genetic Perturbations collection* (*c2.cgp.v2025.1.Hs.symbols.gmt*). The analysis was conducted with 1,000 permutations (45). Enrichment analyses of transcription factors and pathways in cell line clusters were performed using differentially expressed genes in Enrichr tool (48). Visualization of GSEA and Enrichr results was carried out using SRplot (44), as bubble and bar plots, respectively.

### Statistics

Statistical analysis was performed using GraphPad Prism version 9 software. The percentage of tumor-free mice at the endpoint was assessed using Kaplan–Meier survival curves and the log-rank (Mantel–Cox) test. The incidence of tumor dissemination and differences in histological tumor features were evaluated using contingency tables and Fisher’s exact test. Variations in histological morphology patterns were assessed using the Chi-square test. Details of the statistical analysis of transcriptomic data to identify differentially expressed genes are provided in the *Supplemental material*.

For all other experiments, the normality of data distribution (p-value > 0.05) was tested using the Shapiro–Wilk test. Quantitative data not following a normal distribution were analyzed using non-parametric tests (two-tailed Mann–Whitney U test for two-group comparisons or Kruskal– Wallis test followed by Dunn’s post-hoc multiple comparisons test for multiple-group comparisons). Data with a normal distribution were analyzed using parametric tests (two-tailed unpaired t-test with Welch’s correction for two-group comparisons or ANOVA test followed by Tukey’s post-hoc multiple comparisons test for comparisons involving more than two groups). The number of replicates for each *in vitro* assay is detailed in the respective methods section. The number of animals used in *in vivo* experiments is indicated in the figures and/or Supplemental tables.

## Supporting information

This file contains: Supplemental methods, Supplemental Figures 1-13, Supplemental Tables 1-6

## Ethics Statement

All animal studies were approved by the Institutional Animal Care and Use Committee of Centro de Investigaciones Energéticas, Medioambientales y Tecnológicas (CIEMAT) and the competent regional authority (Animal Welfare Department of the Comunidad de Madrid; PROEX 088/15 and 150.8/21), in compliance with Directive 2010/63/EU and Spanish national regulations (RD 53/2013) for the protection of animals used for scientific purposes. All procedures were conducted in accordance with institutional guidelines to minimize animal suffering. Cages were randomized to experimental groups to minimize environmental bias, and all observers were blinded to the experimental conditions.

## Data availability

All transcriptomic data generated in this study are publicly available in the NCBI’s Gene Expression Omnibus (GEO) repository. Microarrays data from mouse bladder tumors derived from GEMMS have been deposited under accession number GSE295920, while RNA-seq data from mouse tumor cell lines are available under accession number GSE302079. Normalized RNA-seq gene expression data from the TCGA BLCA cohort were downloaded from the UCSC Xena platform (https://gdc.xenahubs.net), specifically from the GDC TCGA BLCA dataset (version 05-09-2024). Transcriptomic data from human BC cell lines were obtained from Supplemental Table 8 of Earl J. *et al* (2015) (34), and are also publicly available under accession numbers GSE5845 and GSE64279. Figure 1A, Supplemental Figure S8A, and Supplemental Figure S13A were partially created with BioRender.com. The data generated in this study are available upon request from the corresponding author.

## AUTHOR CONTRIBUTIONS

EMM, MPE, CR, CS, VGM, MMF, MD, JMP and CSC conceptualized the study and designed the experiments. EMM, MPE, CR, CS, IL, SPN, AMB, IAR, EM, LM, VGM, MMF and CSC conducted experiments, and EMM, MPE, CS, IL, SPN, AMB, IAR, EM, LM, VGM, MMF and CSC acquired data. EMM, MPE, CR, CS, VGM, MMF and CSC analyzed data. EMM, MPE, JMP and CSC wrote and edited the manuscript. EMM and MPE contributed equally to this work and the first author was determined alphabetically.

## ACKNOWLEDGEMENTS

We thank the Histology Laboratory from CIEMAT, namely Pilar Hernandez Lorenzo, for the histological processing of tumor samples and the Laboratory of Cytometry and Cellular Separation, specifically Rebeca Sánchez-Domínguez and Omaira Alberquilla for their help with the flow cytometry protocols and analyzes. We also acknowledge Dr. FX Real (CNIO Madrid, Spain) for critical reading the manuscript and important suggestions. The authors acknowledge the assistance of ChatGPT (OpenAI) in revising the grammar and language of the manuscript.

This study was co-funded by grants from the European Regional Development Fund for Science and Innovation (SAF2015-66015-R and PID2019-110758RB-I00 and PID2023-147517OB-I00 to JMP), Instituto de Salud Carlos III (CIBERONC no. CB16/12/00228) to JMP. Project “FINANCED BY NEXTGENERATION EU FUNDS, WHICH FINANCE THE ACTIONS OF THE RECOVERY AND RESILIENCE MECHANISM (MRR)”. Funding entity: CARLOS III HEALTH INSTITUTE (ISCIII). ISCIII project code: AC22/00015 - Title: “Circulating tumor microenvironment components as predictors of response to immunotherapy in urothelial cancer” to MD. Project also funded by the Scientific Foundation of the Spanish Association Against Cancer (FCAECC) with project ID TRNSC213883DUEN, Transcan-3 JTC2022 and Fundación Eugenio Rodríguez Pascual (FERP-2022- 79 to CR).

EMM was supported by a predoctoral fellowship (BES- 2016-078288) linked to the SAF2015- 66015R grant. MPE was supported by a predoctoral fellowship (PRE2020-093423) associated with the PID2019-110758RB-I00 grant. IL was supported by a predoctoral fellowship from AECC (Spanish Ass. against Cancer), Predoctoral AECC 2019 grant number PRDMA19024LODE. SPN. was supported by an FCT-Fundação para a Ciência e Tecnologia Grant (SFRH/BD/144241/2019). VGM was supported by fellowship INVES222946GARC funded by Fundación Científica de la Asociación Española Contra el Cáncer. LM was supported by fellowship POSTD19036MORA, funded by Fundación Científica de la Asociación Española Contra el Cáncer.

