## Supplementary material for "Cell-of-origin and genetic drivers define advanced bladder cancer subtypes and potential therapeutic response in mouse models": This file contains: Supplemental methods, Supplemental Figures 1-13, Supplemental Tables 1-6

1. Cellular and Molecular Oncology and Genitourinary Tumor Group. Institute of Biomedical Research, Hospital Universitario 12 de Octubre, Madrid, Spain.
2. Molecular and Translational Oncology Division. Centro de Investigaciones Energéticas, Medioambientales y Tecnológicas (CIEMAT), Madrid, Spain.
3. Centro de Investigación Biomédica en Red de Cáncer (CIBERONC), Madrid, Spain.
4. Cancer Biology and Epigenetics Group, Research Center of IPO Porto (CI-IPOP)/CI-IPOP@RISE (Health Research Network), Portuguese Oncology Institute of Porto (IPO-Porto)/Porto Comprehensive Cancer Center Raquel Seruca (Porto.CCC), Porto, Portugal.

**Authorship note:** EMM and MPE contributed equally to this work (See author contributions). JMP and CSC are co–corresponding authors.

### SUPPLEMENTAL MATERIAL

**This file contains:**

**Supplemental methods**

**Supplemental Figures 1-13**

**Supplemental Tables 1-6**

#### Supplementary material and methods

##### Cloning of the mouse *Krt20* promoter

To clone the regulatory elements of the murine *Krt20* gene, two upstream sequences—1 kb and 1.5 kb from the transcription start site—were selected (**Supplemental Figure S1A**). These DNA fragments were amplified from murine genomic DNA (C57BL/6J × DBA/2J background) using nested PCR (primer sequences are provided in the table below). The PCR products were cloned into the pGL3 vector using *NheI* and *HindIII* restriction sites. Resulting plasmids were transformed into XL-Blue competent *E. coli* and cultured overnight at 37 °C on LB-agar plates (Luria-Broth, Pronadisa) containing ampicillin (100 µg/mL) for selection. Plasmid DNA was extracted from selected bacterial colonies using the NucleoBond Xtra Midi kit (Macherey-Nagel), followed by a phenol/chloroform purification step to ensure DNA quality. Plasmid constructs were initially screened by restriction digestion with *NheI* and *HindIII*. Selected positive clones were further verified by Sanger sequencing (STAB VIDA, Lisbon, Portugal) using the primers listed in the table below.

| Primer Name | Primer Sequence (5'-3') |
|---|---|
| K20 Forward-1.5Kb | ATATCAGCAGCTTCTGGGAAGTTAGATGTGA |
| K20 Forward-1.5Kb-**NheI** | GGC**GCTAGC**ATATCAGCAGCTTCTGGGAAGTTAG |
| K20 Forward-1Kb | ACTAATTAGATCCAGTTGGTTATTTTTAAAAAGAA |
| K20 Forward-1Kb-**NheI** | GGC**GCTAGC**ACTAATTAGATCCAGTTGGTTATT |
| K20 Reverse | CATCTGGGATGTAGGGAGGCAACGCCTGTA |
| K20 Reverse-**HindIII** | GGC**AAGCTT**CATCTGGGATGTAGGGAGGCAACG |
| pGL3 Fw 2 | CTTTATGTTTTTGGCGTCTTCCA |
| pGL3 Rev 3 | CTAGCAAAATAGGCTGTCCC |

##### *In vitro* validation of the mouse *Krt20* promoter

First, human bladder cancer (BC) cell lines were assessed for keratin 20 expression by immunofluorescence. The J82 BC cell line was used as a positive control due to its high keratin 20 expression, while the 253J cell line served as a negative or low-expression control (**Supplemental Figure S1B**). These cell lines were transfected with either empty or previously constructed pGL3 vectors, as well as the psiCheck vector, using FuGENE-6 (Roche) according to the manufacturer's instructions. Given that the pGL3 vector encodes the *firefly* luciferase gene, the regulatory activity of the cloned *Krt20* promoter sequences was assessed using Dual-Luciferase Reporter Assays (Promega), following the manufacturer's protocol (**Supplemental Figure S1C**). Based on the *in vitro* results, the 1.5 kb fragment was selected as the most suitable carrier of *Krt20* regulatory elements.

**Histological, immunohistochemical, and immunofluorescence analyses**

Tissue samples were fixed in formalin and embedded in paraffin (FFPE), then stained with hematoxylin and eosin (H&E) following standard protocols. FFPE tissue sections were processed for immunohistochemistry (IHC), including deparaffinization, antigen retrieval in sodium citrate buffer, and blocking with horse serum. Samples were incubated overnight at 4 °C with primary antibodies (**Supplemental Table S4**), followed by incubation with secondary antibodies and signal amplification using a biotin–avidin–peroxidase system (**Supplemental Table S4**). Staining was visualized using DAB, counterstained with hematoxylin, and mounted with DPX. Immunofluorescence (IF) analysis was performed on both fixed cultured cells and deparaffinized tissue sections. The protocol included permeabilization, blocking, and incubation with primary and fluorophore-conjugated secondary antibodies (**Supplemental Table S4**). Samples were mounted with Mowiol containing DAPI, and images were acquired using a Leica DM2000 LED microscope for IHC and a Zeiss Axio Imager microscope for IF.

The presence of metastases, tumor morphology, and histological characteristics were evaluated by a veterinary pathologist. Necrosis, immune cell infiltration, and pleomorphic giant cells were recorded when they represented a significant proportion of the tumor. Bone metaplasia and myxoid stroma were assessed qualitatively as present or absent.

**Flow cytometry analysis**

Cells were seeded at a concentration of 10,000 cells/mL in a final volume of 10 mL in p100 dishes and treated with their respective $IC_{50}$ of palbociclib or 0.01% DMSO for 48 hours. For cell cycle analysis, cells were collected along with their growth and wash media, centrifuged, and fixed in 70% ethanol overnight at 4°C. The next day, samples were treated with RNase and stained with propidium iodide. Apoptosis and PD-L1 expression analysis followed a similar procedure, with cells collected, washed, and incubated with FcR blocking reagent before staining. For apoptosis assessment, cells were stained with Annexin V (2 µL; Ref 556421 BD Biosciences) and DAPI (1 µL; Roche), defining populations as live (Annexin V-, DAPI-), early apoptotic (Annexin V+, DAPI-), late apoptotic (Annexin V+, DAPI+), or necrotic (Annexin V-, DAPI+). For PD-L1 expression, cells were additionally stained with an anti-PD-L1 antibody (**Supplemental Table S4**) and analyzed exclusively in the live cell population. All experiments were conducted at least three times, with three technical replicates per condition.

For immune population analysis in *in vivo* experiments, disaggregated tissues were incubated with FcR blocking reagent, stained with viability dyes and specific antibodies (**Supplemental Table S4**), and resuspended in FACS buffer before flow cytometry analysis.

In all experiments, data acquisition was performed using a Becton Dickinson LSR Fortessa analyzer, and results were analyzed with FlowJo 7.6.5 software.

**Cell viability assays**

Cells were seeded at a density of 1,000 cells or 1,500 cells (in the case of cells obtained from QKO and DKO mice, respectively) per well in 96-well plates in quintuplicate. Before adding the compounds, cells were allowed to attach to the bottom of the wells for 24 hours. Then, cells were incubated with 100 µl drug-supplemented medium (using working solutions described in *Material and methods* section) or treated with DMSO (vehicle) at 0.01%. After drug incubation for 48 hours, cell viability was analyzed with XTT Cell Proliferation Kit II (Roche) according to the manufacturer's instructions and absorbance was measured at 490 nm using a Genios Pro microplate reader (Tecan). The half-maximal inhibitory concentration ($IC_{50}$) values were determined using Prism 9 (GraphPad Software v9.0). Each experiment was performed three times.

**RNA extraction**

Total RNA was extracted from FFPE tissue sections using the miRNeasy FFPE kit (Qiagen), or from fresh tissue/cells using the RNeasy Mini kit (Qiagen) following the manufacturer's instructions. The total RNA concentration was quantified using a NanoDrop One spectrophotometer (Controltecnica) for RT-qPCR, and with a Qubit fluorometer for RNA-seq analysis.

**RT-qPCR**

1 µg of total ARN was reverse transcribed with the High-Capacity cDNA Reverse Transcription kit (Applied Biosystems) according to manufacturer's protocol. cDNA was diluted 20 times and 4 µL of this diluted cDNA per reaction was subjected to quantitative qPCR using GoTaq PCR Master Mix (Promega) and both specific primers (**Supplemental Table S5**) at a final concentration of 0.5 µM. qPCRs were run on a QuantStudio™ 6 Flex Real-Time PCR (Thermo Fisher Scientific, Waltham, US) with the standard amplification program (50°C for 2 min and 90°C for 10 minutes steps, 40 consecutive cycles of 95°C/15 seconds and 60°C/1 min). Melting curves were performed to verify primer specificity and dimers absence. The efficiency of the reaction was calculated for each pair of primers by serial dilutions. Results were normalized to mouse *Tbp*/*GusB* expression, and relative expression was calculated using ΔΔCt method.

**DNA extraction and genotyping by PCR**

Cell pellets were lysed using a lysis buffer containing 100 mM Tris-HCl (pH 8.5), 5 mM EDTA, 200 mM NaCl, 0.2% SDS, and 500 µg/mL proteinase K, and incubated at 55°C for at least 2 hours. DNA was then precipitated with isopropanol and washed with 70% ethanol. Finally, the DNA was resuspended in 1X TE buffer. Genotyping of the cell lines was performed by PCR using 1 µL of DNA from each sample, 5 µL of GoTaq Green Master Mix DNA polymerase (Promega), 0.5 µL of specific primer pairs for each gene (**Supplemental Table S6**), and 3 µL of water. In all cases, the following standard amplification program was used: 95°C for 10 minutes, followed by 35 consecutive cycles of 95°C for 15 seconds, 58°C for 15 seconds, and 72°C for 50 seconds, with a final extension step at 72°C for 7 minutes.

### Supplementary figures:

#### Supplemental Figure 1

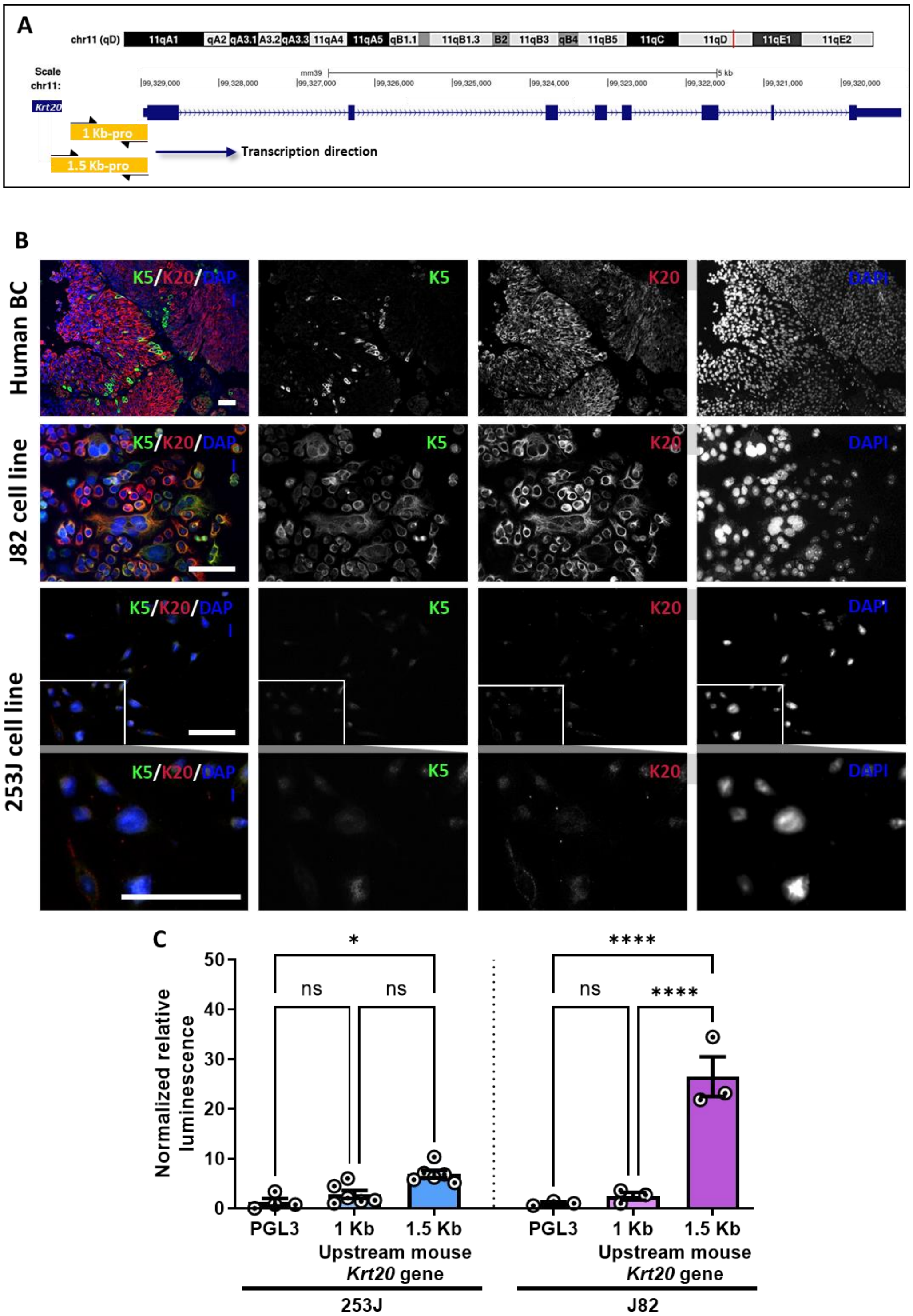


**Supplementary Figure S1. Cloning and *in vitro* characterization of murine *Krt20* promoter. A.** Selected sequences of 1 Kb and 1.5 Kb (shown as yellow rectangles) upstream of the start codon of the murine *Krt20* gene (located in Ch11qD; marked with a red line). The scheme of the mouse *Krt20* gene is shown; the thin parts of the element represent untranslated regions (UTRs), whereas the thick parts represent exons. The scale is in kilobase pairs (Kb). **B.** Immunofluorescence assays to evaluate the expression of keratins K5 (green) and K20 (red) in a human BC tissue and in human BC cell lines. Nuclei are stained with DAPI (blue). Scale bars = 200 µm. In the last row, a high magnification of 253J cells is shown. **C.** Luciferase assay results to evaluate the promoter functionality of the cloned sequences (represented in figure A) in the PGL3 plasmid in both 253J and J82 cell lines. Mean ± SEM and individual values are shown. ns = not significant; *p-value< 0.05; ****p-value< 0.0001.

#### Supplemental Figure 2

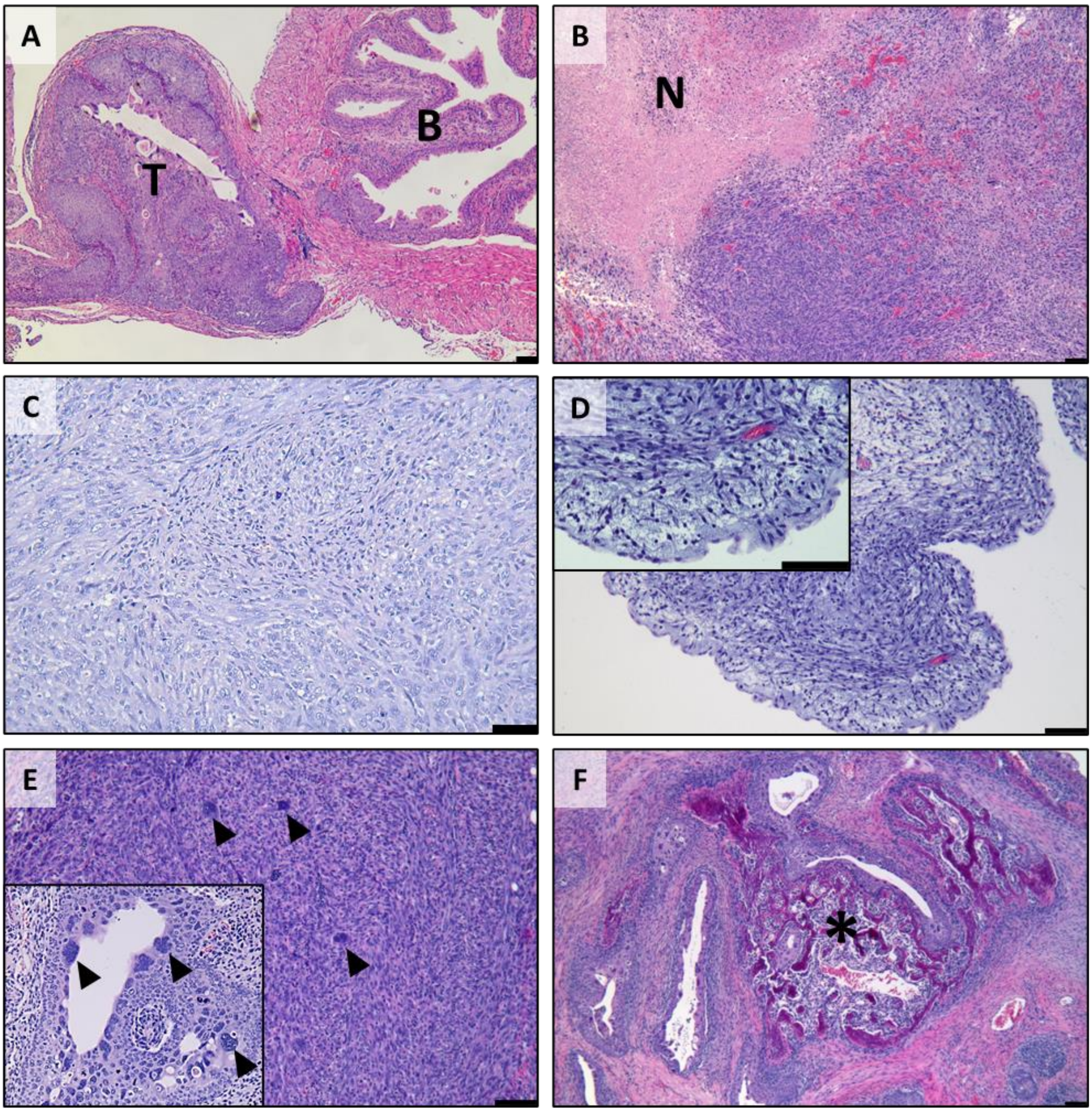


**Supplementary Figure S2. Histological characteristics of tumors from the different BC mouse models. A.** Invasive tumor arising from the bladder in a QKO mouse. **B-F.** Representative H&E staining showing the most common histological features of tumors in the various models, including necrosis (B), immune cell infiltration (C), myxoid stroma (D), giant pleomorphic cells (E) and bone metaplasia with calcification (F). In D, a high magnification is shown. In E, the insert shows giant pleomorphic cells also in a differentiated tumor. Scale bars = 200 µm. N = necrosis; head arrows = giant pleomorphic cells; * = bone metaplasia; T = tumor; B = bladder.

#### Supplemental Figure 3

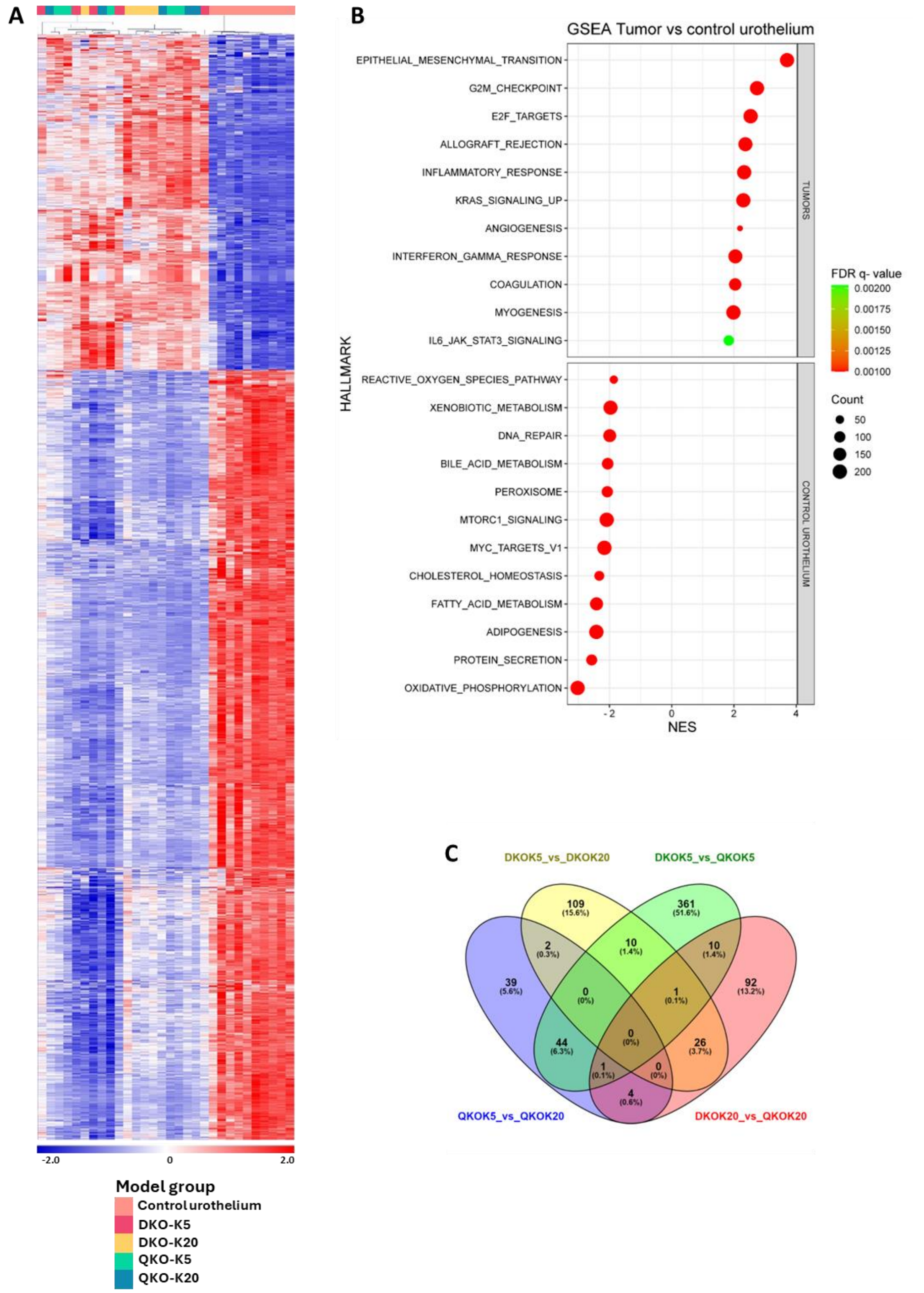


**Supplementary Figure S3. Transcriptomic analysis comparing tumors from the different mouse models with normal urothelium. A.** Heatmap illustrating the clustering of differentially expressed (DE) genes between control tissue and tumors (fold change ±3 and FDR p-value < 0.00005). **B.** Gene Set Enrichment Analysis (GSEA) identifying the most significantly enriched gene sets from the cancer hallmark molecular signature when comparing tumors (top panels) and control urothelium (bottom panels). Normalized enrichment score (NES), FDR q-value, and gene set size (count) are provided. **C.** Venn diagram showing the overlap of DE genes across the different BC mouse models comparisons (fold change ±2 and p-value < 0.05).

#### Supplemental Figure 4

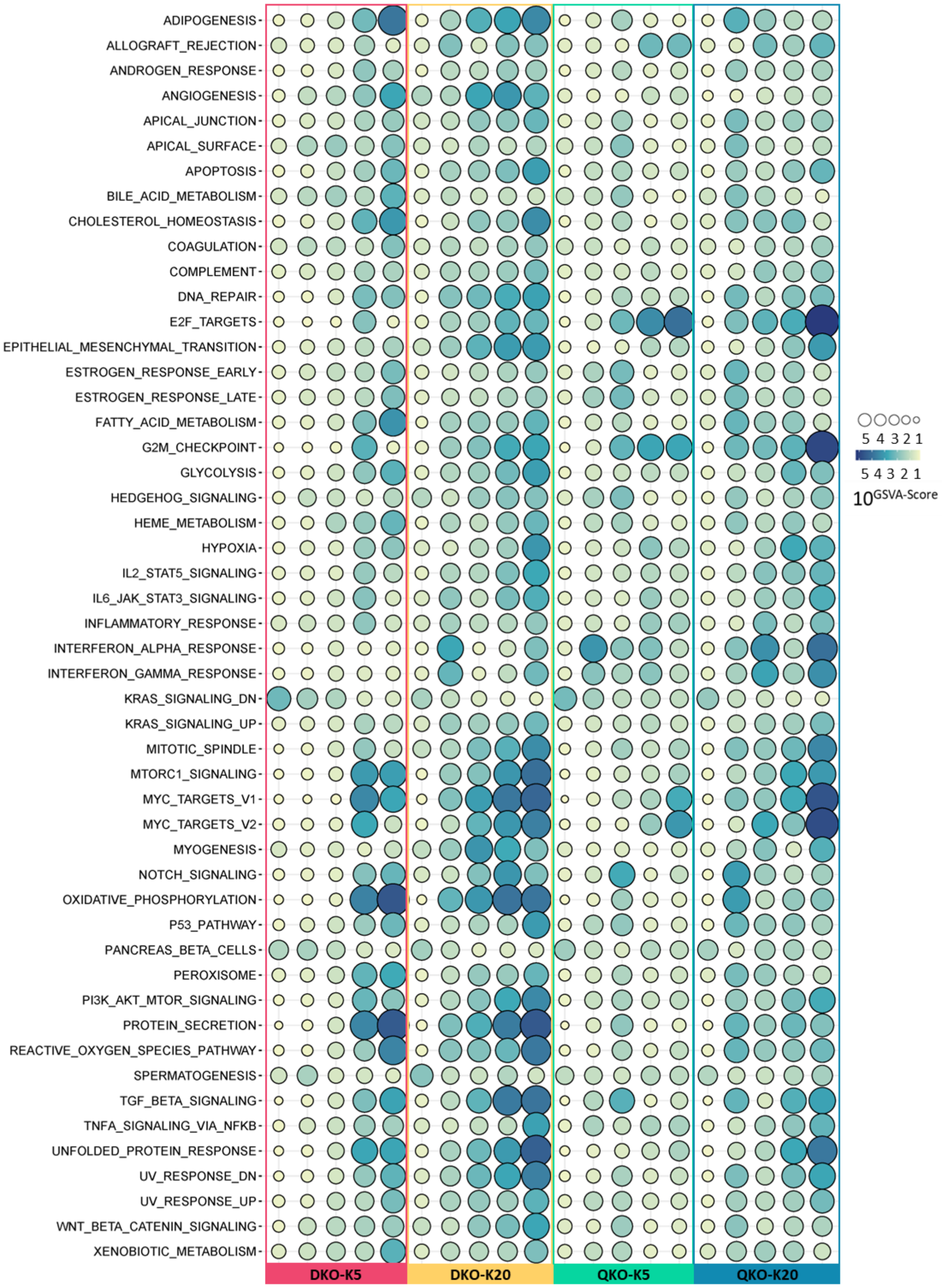


**Supplementary Figure S4. Gene Set Variation Analysis (GSVA) scores for tumors across models illustrating all gene sets from the cancer hallmark molecular signature.** Scores are presented as $10^{\text{GSVA-score}}$.

#### Supplemental Figure 5

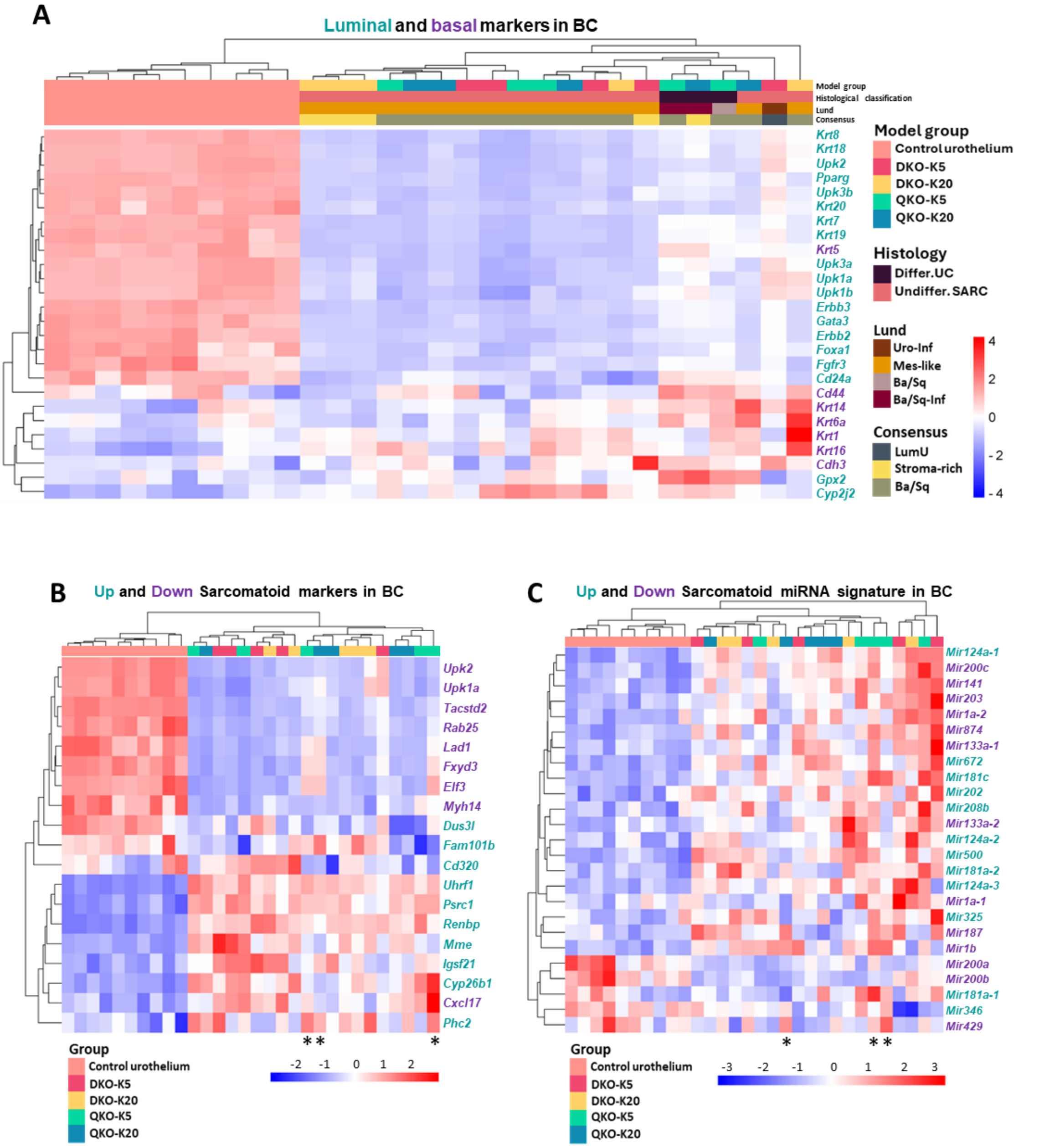


**Supplementary Figure S5. Clustering of tumors from the different mouse models according to BC gene signatures. A-C.** Heatmaps displaying hierarchical clustering of tumors across mouse models based on: (A) luminal and basal markers of BC, (B) mRNA and (C) miRNA sarcomatoid signature in BC. In A, tumor morphology and molecular classification associations are indicated. Luminal markers are highlighted in blue, while basal markers are shown in purple. In B and C, asterisks denote differentiated urothelial carcinoma; upregulated genes are highlighted in blue, while downregulated genes are shown in purple.

#### Supplemental Figure 6

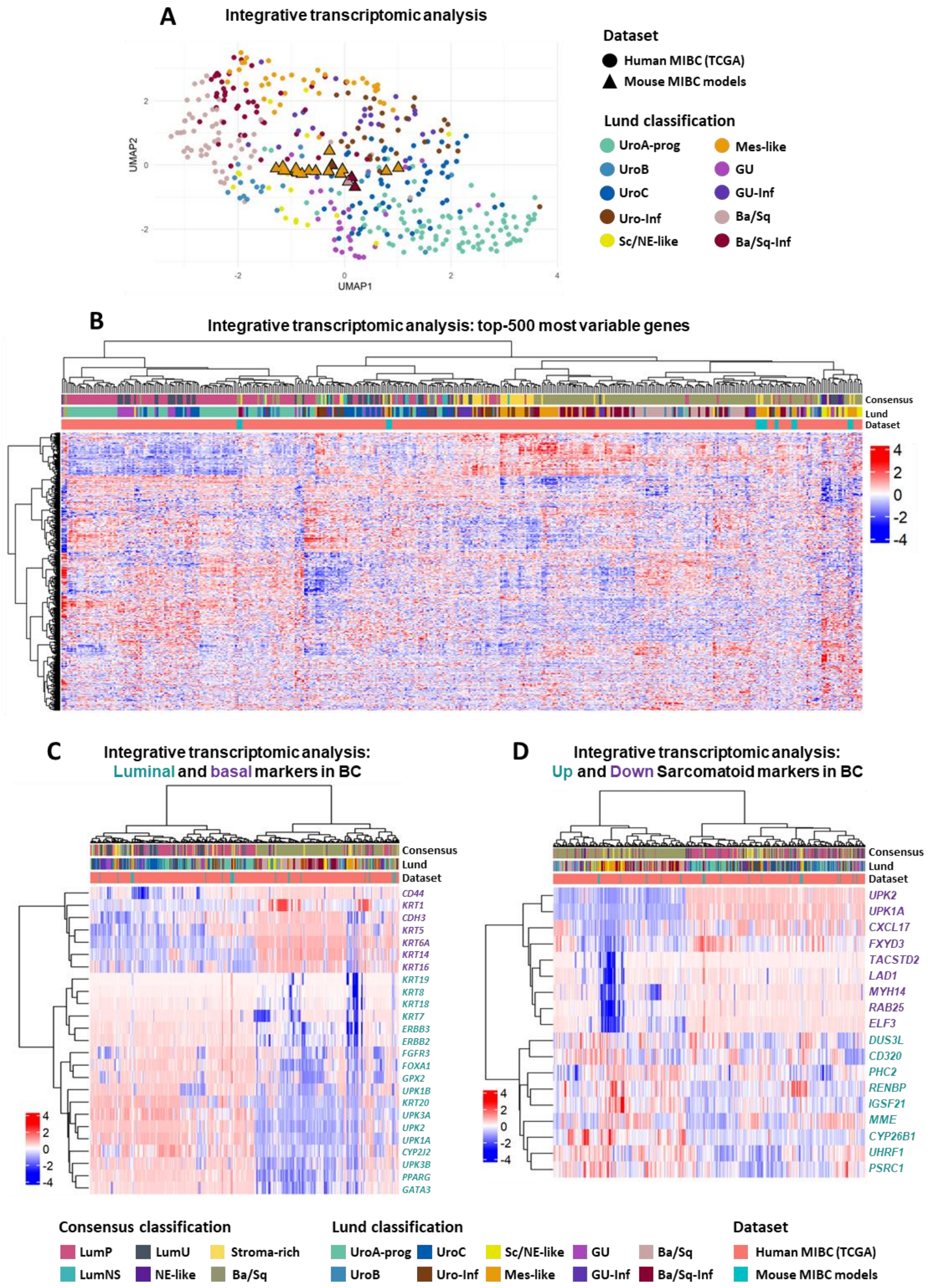


**Supplementary Figure S6. Integrative molecular clustering of mouse BC models and TCGA MIBC samples. A.** UMAP plot illustrating integrative transcriptomic analysis of tumors from BC mouse models together with MIBC samples from TCGA database, with tumor distribution colored according to the Lund molecular classification. **B-D.** Heatmaps displaying hierarchical clustering of mouse and human tumors based on: (B) the 500 most variable genes, (C) luminal and basal BC signature markers, and (D) a sarcomatoid gene expression signature. Consensus and Lund subtype annotations are included. In C, luminal markers are highlighted in blue, while basal markers are shown in purple. In D, upregulated genes are highlighted in blue, while downregulated genes are shown in purple.

#### Supplemental Figure 7

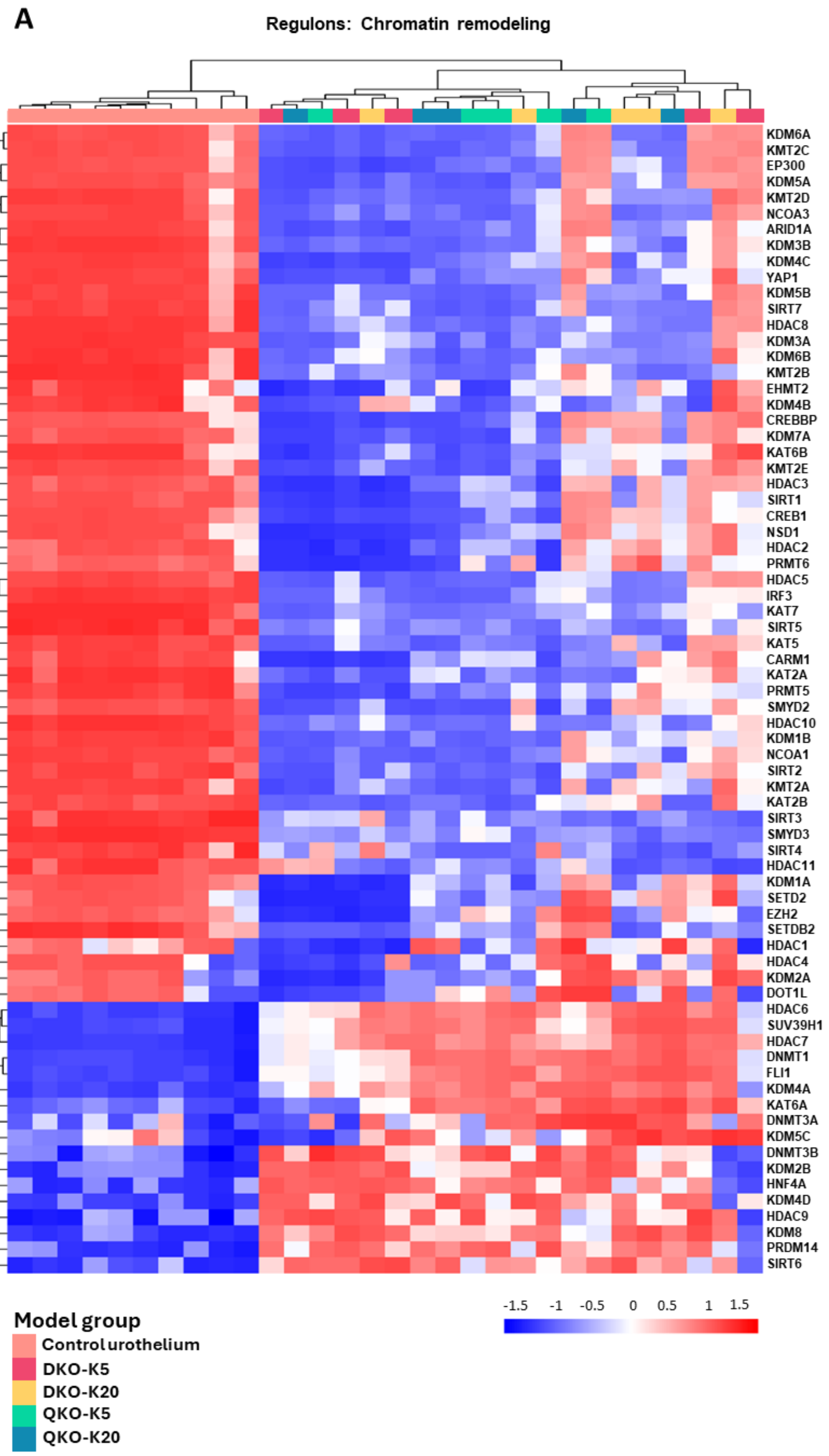


**Supplementary Figure S7. Analysis of chromatin remodeling regulons. A.** Heatmaps showing hierarchical clustering of tumors across mouse models based on regulon activity of chromatin remodeling associated with BC.

#### Supplemental Figure 8

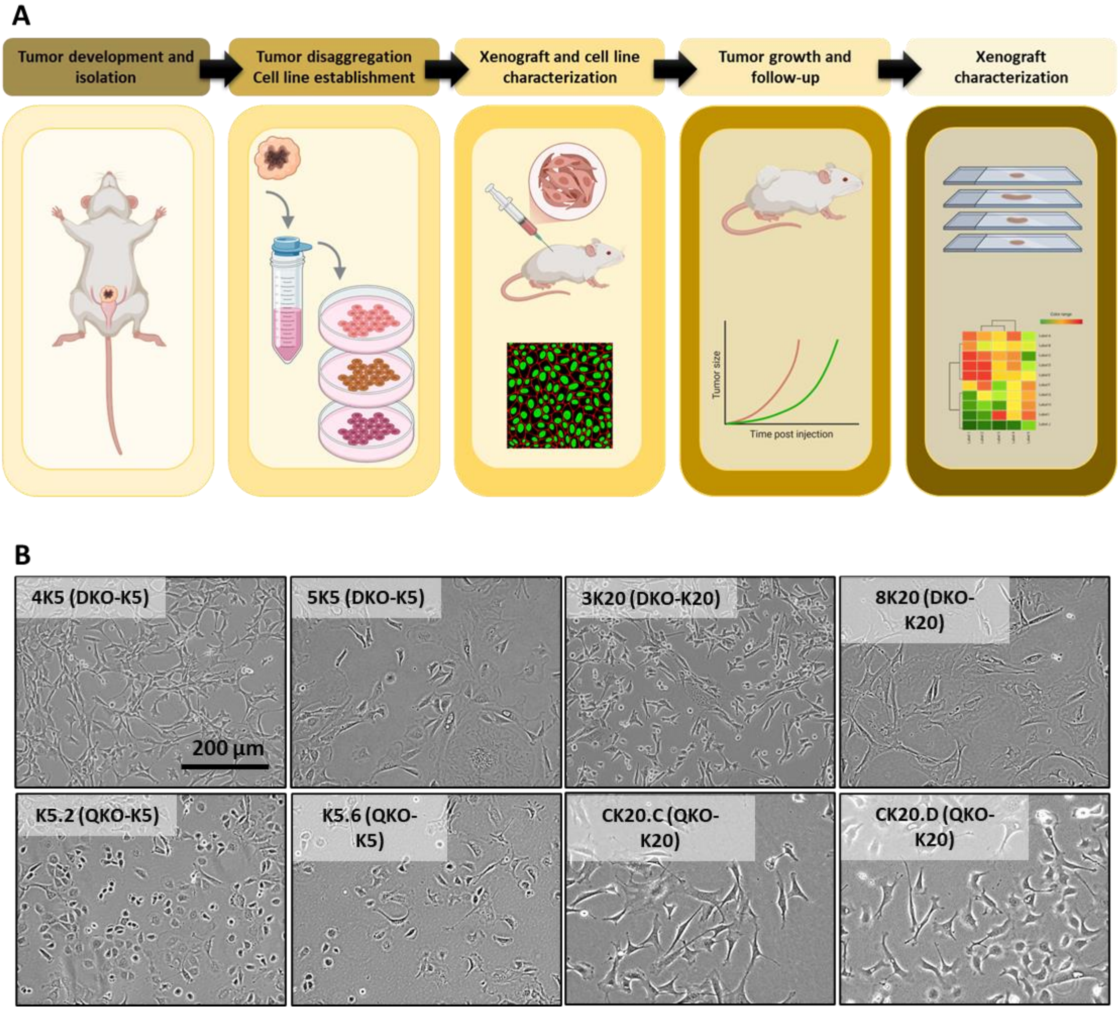


**Supplementary Figure S8. Establishment of tumor cell lines from different models. A.** Schematic representation of tumor isolation and disaggregation to derive tumor cells lines, followed by their characterization and evaluation of growth capacity in immunocompetent models. **B.** Representative images of some established cell lines showing their morphology.

#### Supplemental Figure 9

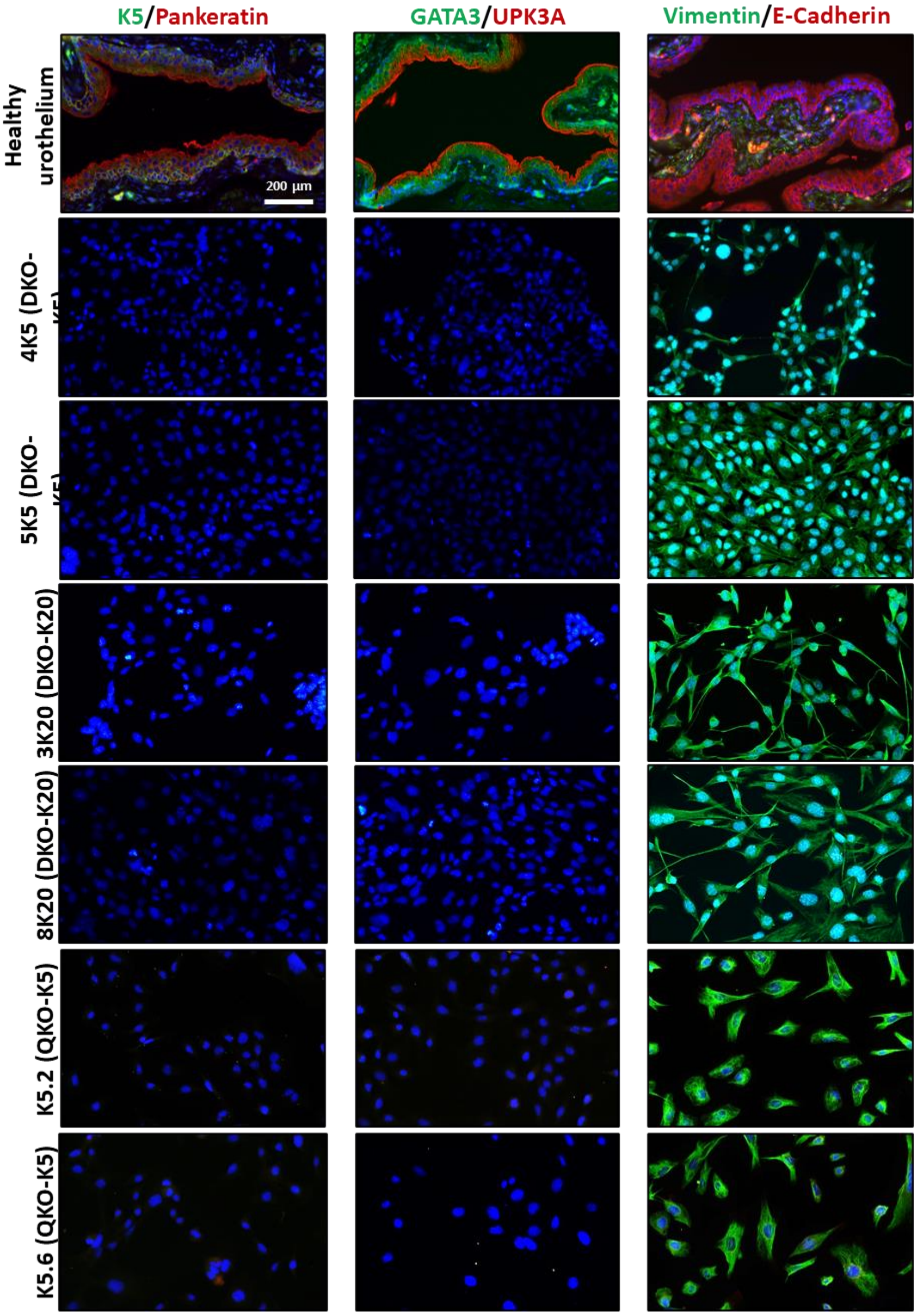


**Supplementary Figure S9. Immunofluorescence analysis of epithelial and mesenchymal biomarkers in various established cell lines.** Healthy urothelium was included as a positive control. Scale bar = 200 µm. All images were acquired using the same magnification and scale settings.

#### Supplemental Figure 10

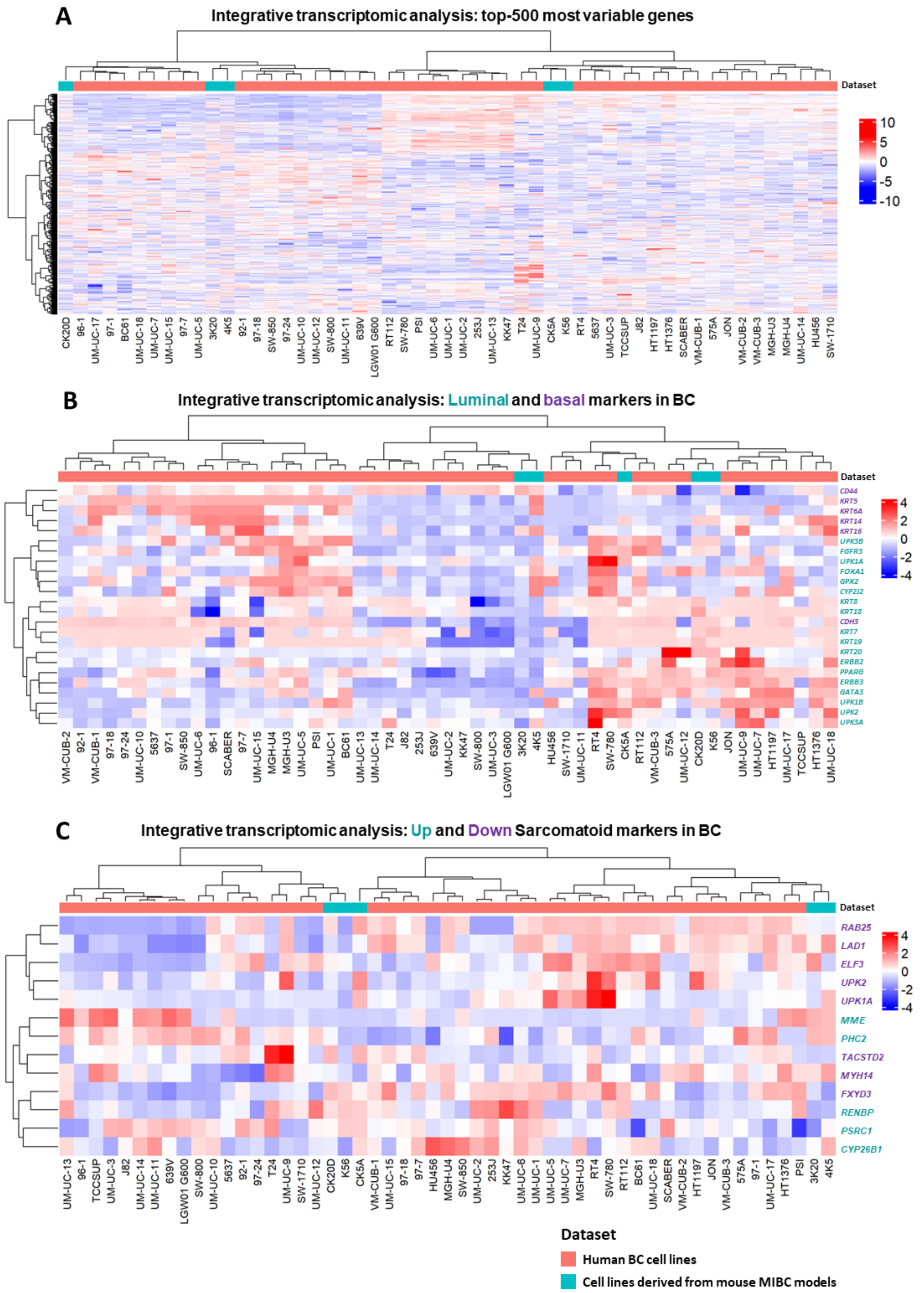


**Supplementary Figure S10. Integrative molecular clustering of mouse BC cell lines and human BC cell lines. A-C.** Heatmaps displaying hierarchical clustering of mouse and human BC cell lines based on: (A) the 500 most variable genes, (B) luminal and basal BC signature markers, and (C) a sarcomatoid gene expression signature. In B, luminal markers are highlighted in blue, while basal markers are shown in purple. In C, upregulated genes are highlighted in blue, while downregulated genes are shown in purple.

#### Supplemental Figure 11

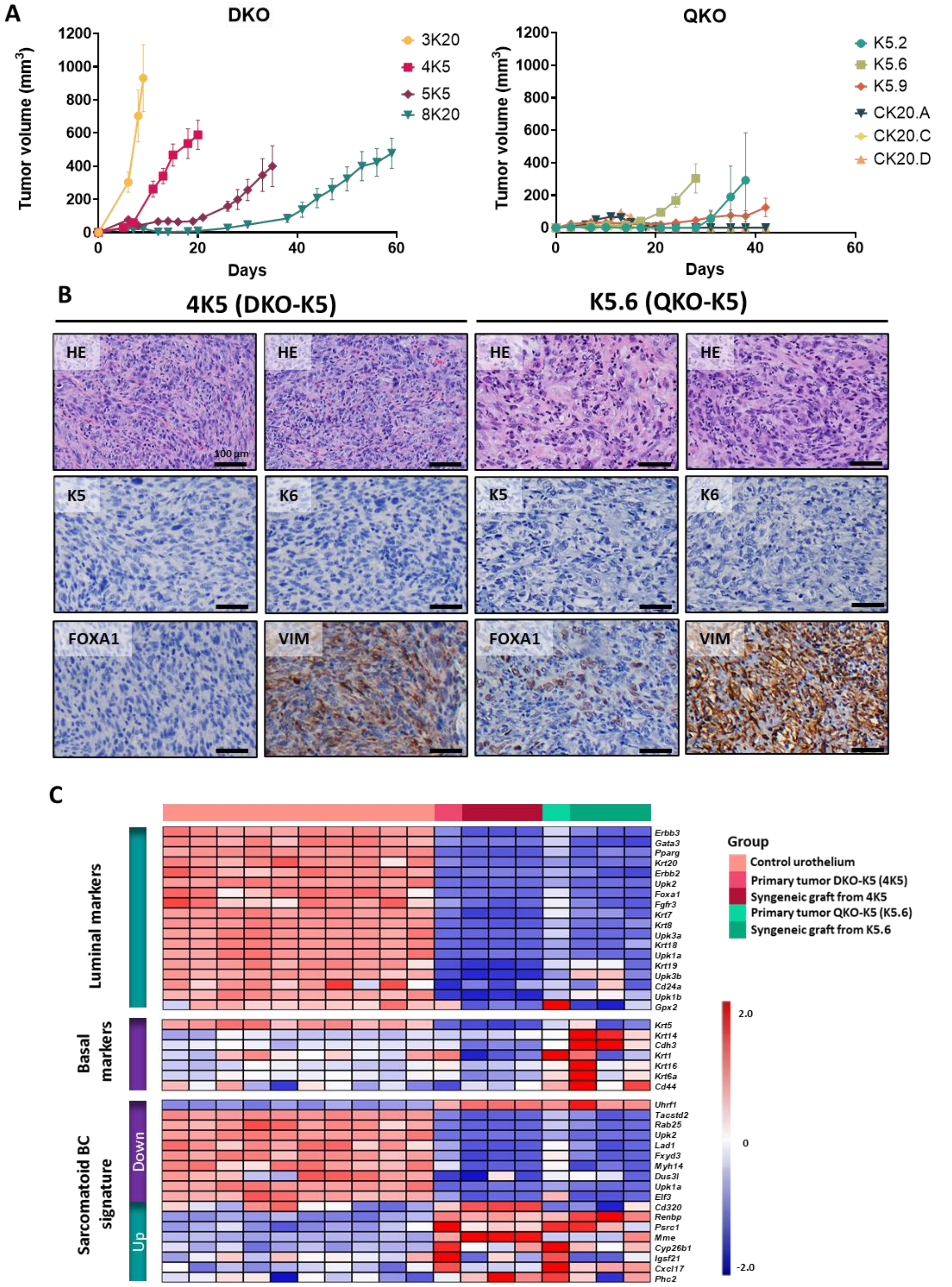


**Supplementary Figure S11. Characterization of immunocompetent syngeneic graft BC mouse models. A.** Tumor growth following heterotopic injection of established cell lines (DKO and QKO) into immunocompetent mice. Data are presented as mean ± SEM. **B**. Representative images of tumors formed after injection of 4K5 (DKO-K5) and K5.6 (QKO-K5), showing tumor morphology (H&E staining) and the expression of luminal (FOXA1), basal (K5/K6), and mesenchymal (vimentin) markers analyzed by immunohistochemistry. **C.** Transcriptomic analysis of bladder tumors derived from immunocompetent syngeneic grafts, compared to their primary tumor of origin, using luminal, basal, and sarcomatoid gene signatures from human BC.

#### Supplemental Figure 12

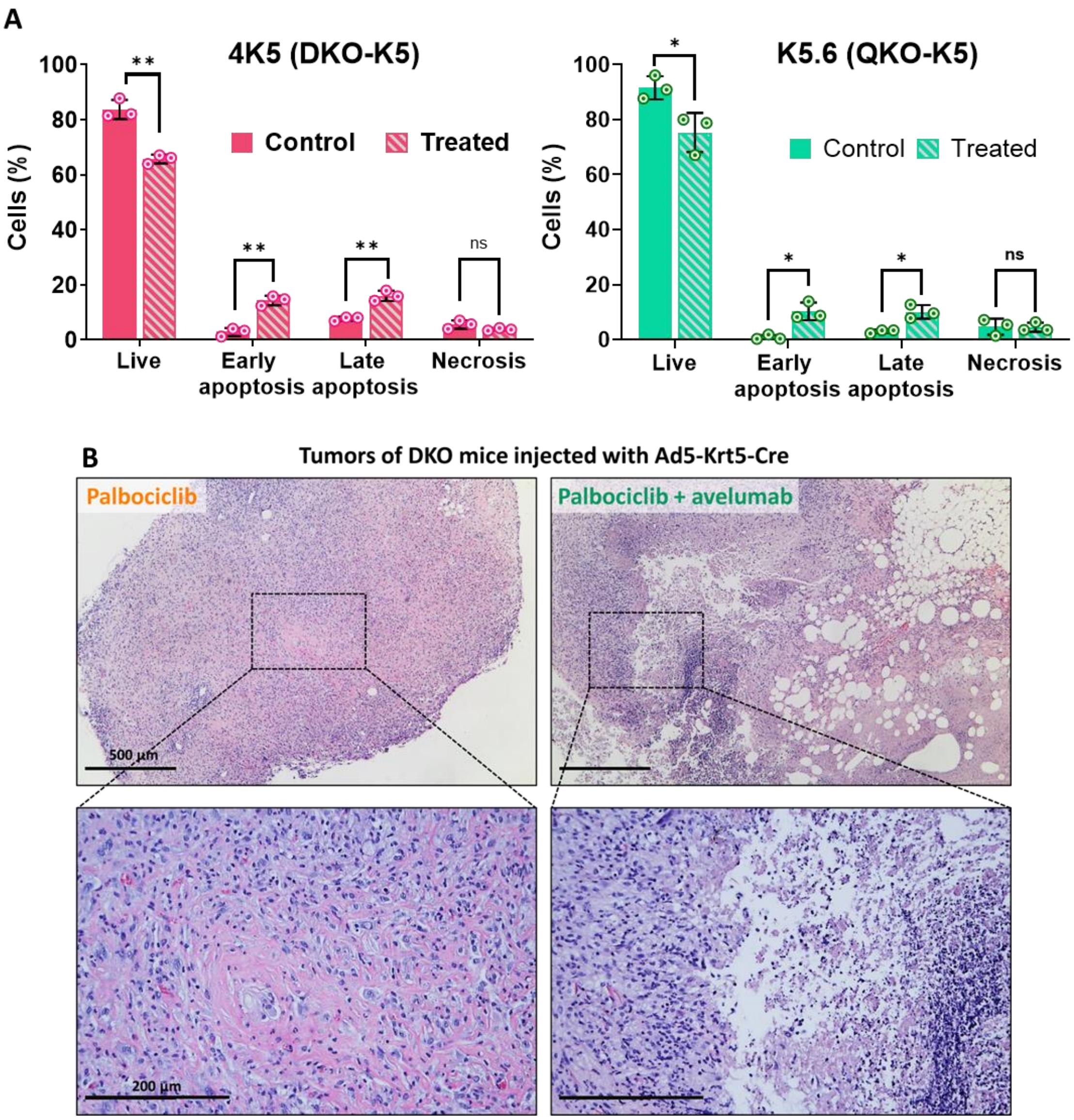


**Supplementary Figure S12.** ***In vitro*** **and** ***in vivo*** **analysis of tumor cell response to treatment. A.** *In vitro* analysis of apoptosis and cell death in mouse BC cell lines derived from K5-positive cells isolated from DKO (4K5; left) and QKO (K5.6; right) tumors following 48-hour treatment with palbociclib at the $IC_{50,}$ analyzed by flow cytometry. Data are presented as mean ± SD from three independent experiments, with the average value of each experiment shown as a single data point. **B.** Proof-of-concept experiment with palbociclib alone (n = 5) or combined with avelumab (n = 5) in DKO mice injected with Ad-K5-Cre into the bladder lumen. A control group of 5 mice was used. Representative H&E-stained tumor sections are shown post-treatment. ns = not significant; *p-value< 0.05; **p-value< 0.01.

#### Supplemental Figure 13

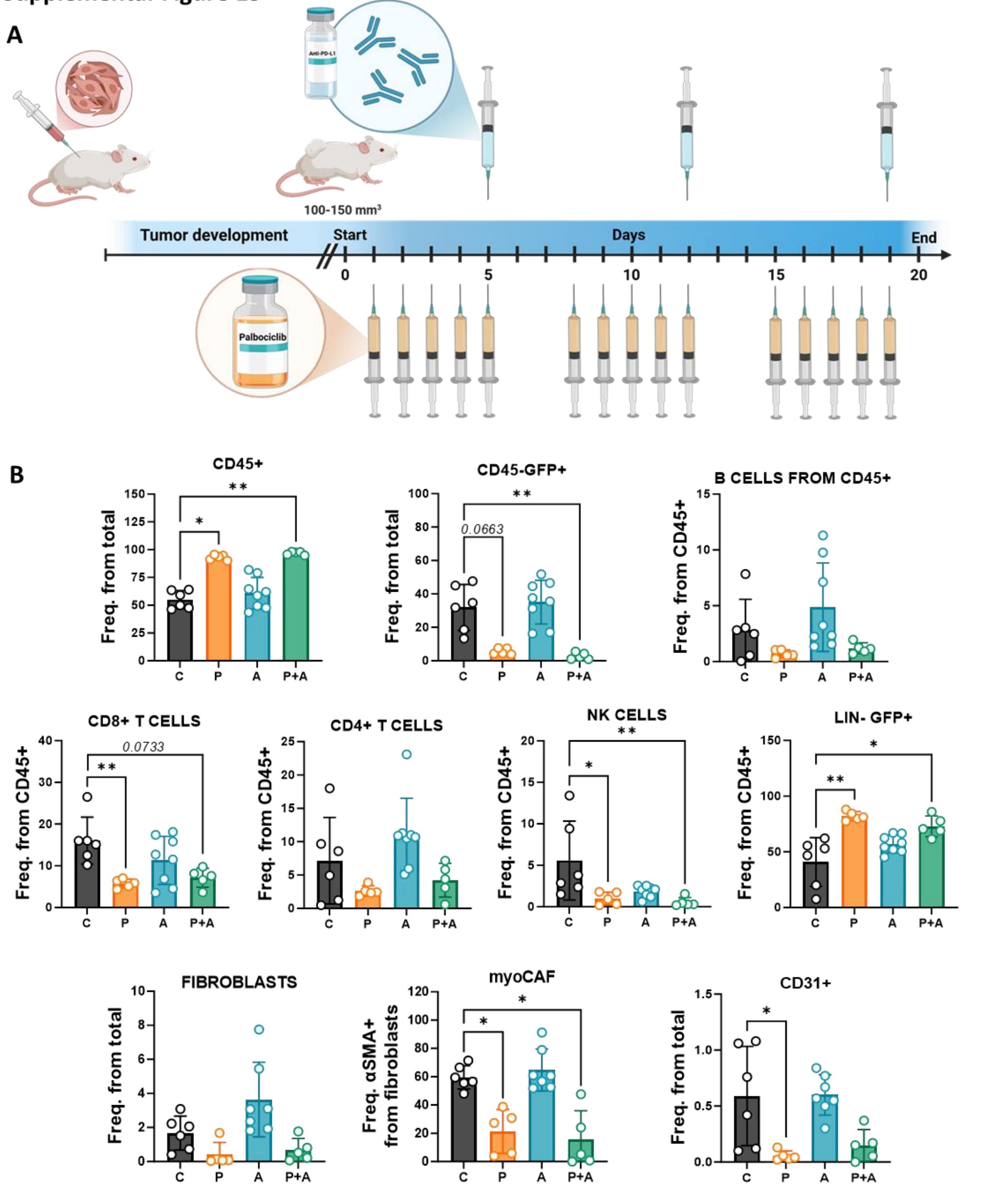


**Supplementary Figure S13. Syngeneic graft BC models to evaluate palbociclib and avelumab combination therapy. A.** Schematic representation of treatment regimens for monotherapy and combination therapy treatments in immunocompetent syngeneic graft BC mouse models. **B.** Analysis of alterations in tumor microenvironment populations of tumor derived from DKO-K5 cells following treatment with monotherapy and combination therapy in heterotopic tumors. Data from each individual are shown, along with the mean ± SD. Only significant differences are shown, along with trends indicated by the p-value. C = control; P = palbociclib; A = avelumab; P+A = combination palbociclib plus avelumab. LIN- = negative lineage. *p-value< 0.05; **p-value< 0.01.

#### Supplementary tables:

| Genotype | Adenovirus | N | Tumor incidence: % Tumor (Tumor/total) | Tumor incidence: Median survival, days (range) | Histopathological phenotype | Dissemination: Carcinomatosis | Dissemination: Distant metastasis |
|---|---|---|---|---|---|---|---|
| DKO (*Pten* $^{flox/flox}$; *Trp53* $^{flox/flox}$) | *Ad5-Krt5-Cre* | 29 | 55.2 (16/29) | 213 (92-280) | Sarcomatoid and invasive urothelial carcinomas (100%) | 62.5% (10/16) | 12.5% (2/16) |
| | *Ad5-Krt20-Cre* | 20 | 90 (18/20) | 138 (89-221) | Sarcomatoid and invasive urothelial carcinomas (100%) | 16.7 % (3/18) | 0% (0/18) |
| QKO (*Pten* $^{flox/flox}$; *Trp53* $^{flox/flox}$; *Rb1* $^{flox/flox}$; *Rbl1* $^{-/-}$) | *Ad5-Krt5-Cre* | 42 | 78.6 (33/42) | 95 (55-154) | Invasive urothelial carcinomas, sarcomatoid (57.6%), differentiated (9.1%) or with mixed pattern (33.3%) | 78.8% (26/33) | 30.3% (10/33) |
| | *Ad5-Krt20-Cre* | 36 | 77.8 (28/36) | 68 (53-88) | Invasive urothelial carcinomas, sarcomatoid (42.9%), differentiated (10.7%) or with mixed pattern (46.4%) | 60.7% (17/28) | 21.4% (6/28) |

**Supplementary Table S1. Overview of four BC mouse models, including tumor incidence, latency, morphology, and dissemination characteristics.**

| | Groups | DKO-K5 | DKO-20 | QKO-K5 | QKO-K20 | K5 vs K20 | | | DKO vs QKO | | |
|---|---|---|---|---|---|---|---|---|---|---|---|
| | Number of injected mice (n) | 29 | 20 | 42 | 36 | K5 | K20 | | DKO | QKO | |
| | Total developed tumors (n) | 16 | 18 | 33 | 28 | 49 | 46 | | 34 | 61 | |
| | Tumor incidence (%) | 55.2 | 90.0 | 78.6 | 77.8 | 69 | 82 | p-value | 69 | 78 | p-value |
| Histological pattern | Undifferentiated, Sarcomatoid (%) | 100 | 100 | 57.6 | 42.9 | 71.4 | 65.2 | | 100.0 | 50.8 | |
| | Differentiated, Urothelial carcinoma (%) | 0 | 0 | 9.1 | 10.7 | 6.1 | 6.5 | ns | 0.0 | 9.8 | **<0.0001** |
| | Mixed (%) | 0 | 0 | 33.3 | 46.4 | 22.4 | 28.3 | | 0.0 | 39.3 | |
| Histological features of tumors | Myxoid stroma (%) | 12.5 | 33.3 | 45.5 | 50.0 | 34.7 | 43.5 | ns | 23.5 | 47.5 | **0.0282** |
| | Giant pleomorphic cells (%) | 31.3 | 88.9 | 27.3 | 53.6 | 28.6 | 67.4 | **0.0002** | 61.8 | 39.3 | ns (0.0532) |
| | Immune cell infiltrate (%) | 18.8 | 11.1 | 72.7 | 67.9 | 55.1 | 45.7 | ns | 14.7 | 70.5 | **<0.0001** |
| | Necrosis (%) | 6.3 | 0 | 33.3 | 50.0 | 24.5 | 30.4 | ns | 2.9 | 41.0 | **<0.0001** |
| | Bone metaplasia (%) | 0 | 0 | 57.6 | 35.7 | 57.6* | 35.7* | **0.0464** | 0.0 | 47.5 | **<0.0001** |

**Supplementary Table S2. Histological characterization of tumors from the four BC mouse models, comparing histological patterns and common features.** n = number of animals or tumors. ns = not significant. * Bone metaplasia percentages are reported only for tumors from QKO mice. P-values near significance are indicated in parentheses.

| Tumor genotype | Cell of origin/promoter | Cell lines | Heterotopic syngeneic graft | | | |
|---|---|---|---|---|---|---|
| | | | Incidence | Regression | End-volume average | Survival (dpi) |
| DKO (*Pten* $^{-/-}$; *Trp53* $^{-/-}$) | Basal cell/Krt5 | 4K5 | 100% (8/8) | 0% | 590 mm$^3$ | 20 |
| | | 5K5 | 100% (8/8) | 0% | 400 mm$^3$ | 35 |
| | Luminal cell/Krt20 | 3K20 | 100% (8/8) | 0% | 930 mm$^3$ | 9 |
| | | 8K20 | 100% (8/8) | 0% | 480 mm$^3$ | 59 |
| QKO (*Pten* $^{-/-}$; *Trp53* $^{-/-}$; *Rb1* $^{-/-}$; *Rbl1* $^{-/-}$) | Basal cell/Krt5 | K5.2 | 37.5% (3/8) | 67% (2/3) | 290 mm$^3$ | 38 |
| | | K5.6 | 100% (8/8) | 0% | 350 mm$^3$ | 28 |
| | | K5.9 | 100% (8/8) | 50% (4/8) | 125 mm$^3$ | 42 |
| | | CK5.A | ND | ND | ND | ND |
| | | CK5.B | ND | ND | ND | ND |
| | Luminal cell/Krt20 | CK20.A | 100% (8/8) | 100% (8/8) | 0 mm$^3$ | 90 |
| | | CK20.C | 75% (6/8) | 100% (6/6) | 0 mm$^3$ | 90 |
| | | CK20.D | 100% (8/8) | 100% (8/8) | 0 mm$^3$ | 90 |
| | | CK20.E | ND | ND | ND | ND |

**Supplementary Table S3. Summary of established tumor-derived cell lines including incidence, percentage of regression, average end volume, and survival.** ND = not determined; dpi = days post-injection.

| IHC and IF primary antibodies | | | | | |
|---|---|---|---|---|---|
| **Epitope** | **Clone** | **Provider** | **Reference** | **Host** | **Dilution** |
| **Keratin 5** | Polyclonal | Biolegend | 905501 | Rabbit | 1/1000 |
| **Keratin 6** | Polyclonal | Biolegend | 905702 | Rabbit | 1/500 |
| **Keratin 20** | Ks20.8 | Dako | 1S777 | Mouse | 1/1000 |
| **Pankeratin** | AE1/AE3 | Abcam | ab27988 | Mouse | 1/100 |
| **FOXA1** | EPR10881 | Abcam | ab170933 | Rabbit | 1/1000 |
| **Vimentin** | EPR3776 | Abcam | ab92547 | Rabbit | 1/250 |
| **GATA3** | Polyclonal | Sigma Aldrich | SAB4501126 | Rabbit | 1/250 |
| **UPK3A** | C-6 | Santa Cruz | sc-166808 | Mouse | 1/200 |
| **E-cadherin** | 36 | BD | 610181 | Mouse | 1/100 |
| **GFP** | Polyclonal | Invitrogen | A-11122 | Rabbit | 1/200 |
| **Secondary antibodies** | | | | | |
| **Epitope** | **Conjugate** | **Provider** | **Reference** | **Host** | **Dilution** |
| **IgG Rabbit** | Biotin | Jackson | 711-065-152 | Donkey | 1/1000 |
| **IgG Mouse** | Biotin | Jackson | 715-065-151 | Donkey | 1/1000 |
| **IgG Rabbit** | Alexa Fluor 488 | Invitrogen | A11034 | Goat | 1/1000 |
| **IgG Mouse** | Alexa Fluor 594 | Invitrogen | A11005 | Goat | 1/1000 |
| **FACS antibodies** | | | | | |
| **Epitope** | **Fluorochrome** | **Clone** | **Provider** | **Reference** | **Vol/100ul** |
| **CD45** | PerCP | 30-F11 | Biolegend | 103130 | 1 |
| **CD3** | APC-Cy7 | 17A2 | Biolegend | 100222 | 1.5 |
| **CD4** | BV711 | RM4-5 | Biolegend | 100550 | 1 |
| **CD8** | PE | 53-6.7 | BD | 553033 | 1 |
| **NK1.1** | APC | PK136 | Biolegend | 108710 | 2 |
| **B220** | PEFire700 | RA3-6B2 | Biolegend | 103280 | 0.75 |
| **CD31** | PE-Cy7 | 390 | Biolegend | 102418 | 1 |
| **PDPN** | AF-647 | P-mab1 | Biolegend | 156203 | 0.5 |
| **CD140a** | BV605 | APA4 | Biolegend | 135916 | 1 |
| **αSMA** | Cy3 | 1A4 | Sigma-Aldrich | C6198 | 0.5 |
| **Zombie Aqua** | - | - | Biolegend | 423102 | 0.67 |
| **PD-L1** | BV711 | MIH5 | BD | 563369 | 2 |

**Supplementary Table S4. Primary and secondary antibodies used in immunohistochemistry, immunofluorescence, and flow cytometry assays.**

| Gene | Primer sequence 5'-3' | |
|---|---|---|
| ***GusB*** | Forward | GAGGATCAACAGTGCCCATT |
| | Reverse | CAGCCTCAAAGGGGAGGT |
| ***Tbp*** | Forward | GGGAGAATCATGGACCAGAA |
| | Reverse | GATGGGAATTCCAGGAGTCA |
| ***Trp53*** | Forward | GCCCATGCTACAGAGGAGTC |
| | Reverse | AGACTGGCCCTTCTTGGTCT |
| ***Rb1*** | Forward | CACGTGTAAATTCTGCTGCAA |
| | Reverse | ACAGGGCAAGGGAGGTAGAT |
| ***Pten*** | Forward | GAAAGGGACGGACTGGTGTA |
| | Reverse | TAGGGCCTCTTGTGCCTTTA |

**Supplementary Table S5. Primers used for quantitative PCR analysis.**

| Gene | Primer sequence 5'-3' | | Product Size(bp) | Genotype |
|---|---|---|---|---|
| ***Rb1*** | Forward<br>Reverse | GGCGTGTGCCATCAATG<br>AACTCAAGGGAGACCTG | 662 | WT |
| | | | 734 | Floxed |
| | | | 298 | Deleted |
| ***Trp53*** | Forward 1 | AAGGGGTATGAGGGACAAGG | 154 | WT |
| | Reverse | GAGACAGGGTCTTGCTATTGT | 307 | Floxed |
| | Forward 2 | AGAAAGGGCGACTGACTGTG | 203 | Deleted |
| ***Pten*** | Forward 1 | AGTGGCATGTTTTGTCTATGGT | 72 | WT |
| | Reverse | ACGAGTCCTCTGAAAAAGCAGT | 118 | Floxed |
| | Forward 2 | TGGGGCTGCAGGAATTCGATA | 141 | Deleted |
| ***Rbl1*** | Forward | GCAACTTTGGTGGCTCTTCAT | 130 | WT |
| | Reverse 1 | CCACAAGAGTTTCGTGAGCG | 78 | Null |

**Supplementary Table S6. Primers used for genotyping PCR analysis.** WT = wild-type; bp = base pairs. Floxed = allele carrying the loxP site(s). Deleted = sequence generated by recombining the loxP sites. Null = knock-out allele with neomycin resistance cassette.
